# Bioinformatic methods for stratification of obese patients and identification of cancer susceptibility biomarkers based on the analysis of the gut microbiome

**DOI:** 10.1101/2022.11.17.516892

**Authors:** Blanca Lacruz-Pleguezuelos, Lara P. Fernández, Ana Ramírez de Molina, Enrique Carrillo de Santa Pau, Laura Judith Marcos-Zambrano

**Affiliations:** Computational Biology Group, Precision Nutrition and Cancer Research Program, IMDEA Food Institute, CEI UAM+CSIC, 28049 Madrid, Spain; Molecular Oncology Group, Precision Nutrition and Cancer Research Program, IMDEA Food Institute, CEI UAM + CSIC, Carretera de Cantoblanco 8, E-28049 Madrid, Spain

**Keywords:** Obesity, metabolically healthy obesity, gut microbiota, colorectal cancer, random forest, patient similarity network

## Abstract

Obesity has an impact on health by increasing the risk of various diseases. However, these risks might also depend on the metabolic health status, as it seems that metabolically healthy obese subjects are under a reduced risk of suffering comorbidities such as colorectal cancer. The gut microbiome has an effect on obesity and metabolic disorders through several integration pathways, making it a potential therapeutic target for these diseases. In this study, we characterized the gut microbiota of 356 obese and non-obese European individuals with different comorbidities associated with obesity. Using approaches based on supervised machine learning and network biology, we found a set of biomarkers of interest for differentiating metabolically healthy from unhealthy subjects. Then, we performed a linear discriminant analysis of effect size on a population of 1593 colorectal cancer, adenoma and control subjects assembled by the COST Action ML4Microbiome to investigate their role in colorectal cancer risk. Four of our biomarkers appeared in both approaches, suggesting their possible role in colorectal cancer development, prognosis and follow up: *Clostridium leptum, Gordonibacter pamelaeae, Eggerthella lenta* and *Collinsella intestinalis*. Further research via longitudinal studies or experimental validation of these microbial species would be necessary to confirm this association.

## Introduction

The global prevalence of obesity has nearly tripled between 1975 and 2016, with more than 1.9 billion overweight and obese adults worldwide [1]. Recent estimates report that 15.5 % of women and 16.5 % of men in the adult Spanish population are obese [2]. An increased body mass index (BMI) represents a risk factor for non-communicable diseases such as type 2 diabetes (T2D), cardiovascular and inflammatory diseases, and different types of cancer [3–6]. Obesity and its comorbidities are estimated to decrease life expectancy by up to 20 years, and hence represent an important burden for health worldwide [6,7].

However, some obesity patients do not present metabolic abnormalities, a phenotype known as metabolically healthy obesity (MHO) [6]. The BMI does not describe adiposity or body composition, and it is often not enough to determine the health status at an individual level. MHO is more often observed in women and its prevalence decreases with age [6]. These patients are characterized by lower levels of ectopic fat storage and seem to be protected against cardiovascular disease (CVD) or T2D when compared to metabolically unhealthy obese (MUO) subjects, an association which has been also observed for colon, breast and gastric cancer [5,6,8]. However, this phenotype is not fully understood yet. The lack of a standardized definition, its stability over time and whether it is a benign condition at all are some of the issues that remain open in the research community [6,9,10].

The gut microbiota (GM) has been suggested as a factor that could influence the development of complex diseases, including obesity and metabolic disorders, in interaction with genetic and environmental factors [11,12]. The GM is involved in processes such as energy homeostasis and immunity, which might explain these associations [11–13]. For instance, energy harvesting by the gut microorganisms is increased in obese and overweight patients, leading to increased adiposity [12]. Moreover, high-fat diets change the GM composition, leading to an increase in circulatory lipopolysaccharides, endotoxemia and low-grade inflammation, which can promote obesity and metabolic disorders [11–13]. Other mechanisms by which the GM has been suggested to affect the development of metabolic disease include the alteration of glucose homeostasis and gut permeability [13].

Thus, a great effort has been made to elucidate the role that the GM plays in the development of obesity and its comorbidities, including T2D or colorectal cancer (CRC). Various studies have tried to discover the mechanisms and the specific microbial signatures involved in these diseases [12–19]. Modulation of the GM by probiotic or prebiotic intake, dietary changes or fecal microbiota transplantation has positive effects on metabolic health and weight loss interventions [20]. Moreover, the GM composition can affect the efficacy of weight loss therapies at an individual level [21]. All this underscores the potential that the GM has in precision medicine.

Recently, some studies have investigated whether the GM composition can differentiate obese and overweight patients with different metabolic phenotypes [15,22,23]. Our aim was to further this research by studying whether the relation between the metabolic health status and the CRC risk is mediated by changes in the GM. With this purpose, we have characterized the GM of 356 obese and lean subjects with different metabolic health status. Then, we have built a random forest model and a patient similarity network to look for patient subgroups as well as for potential biomarkers. In parallel, we performed a differential abundance analysis on a cohort of CRC and adenoma patients, looking for a possible connection between the MUO phenotype and CRC development mediated by the GM.

## Methods

### Data collection

We searched for publicly available whole-genome shotgun sequencing (WGS) or 16S ribosomal RNA gene (16S rRNA) sequencing studies for which metadata regarding the BMI and the metabolic status of the patients were available. We performed a bibliographic search through PubMed using (((obesity) AND (metagenomics)) NOT (mice)) NOT (review[Publication Type]) as search term and reviewed all the results up to the 10th November 2021, as well as the studies referenced by these publications. We also looked for datasets available in the Qiita [24], MGNify [25] and curatedMetagenomicData [26] resources.

Our initial search retrieved 20 studies of interest, with a total of 7277 samples. Out of these studies, only 4 included the necessary metadata for patient classification. Thus, we have used 356 fecal metagenomes from four cohorts: 145 samples from the KarlssonFH_2013 dataset [27] and the 61 control samples from the CRC FengQ_2015 dataset [17], both available in the R package curatedMetagenomicData (version 3.0.10) [26]; and two cohorts from the Metagenomics of the Human Intestinal Tract (MetaHit) project [28]: one formed by 85 healthy patients and another composed of 65 T2D patients [28,29]. Raw reads from these patients were downloaded from the European Nucleotide Archive (ENA) projects PRJEB2054 and PRJEB5224 respectively.

We classified patients following the criteria defined by the BioSHaRE-EU Healthy Obese Project [30]. Briefly, patients were considered metabolically healthy (MH) only if they showed all of the following: low fasting blood glucose (≤ 6.1 mmol/l or ≤ 100 mg/dl), low fasted serum triglycerides (≤ 1.7 mmol/l or ≤ 150 mg/dl), high HDL cholesterol concentrations (> 1.0 mmol/l or > 40 mg/dl in men and > 1.3 mmol/l or > 50 mg/dl in women), systolic blood pressure ≤ 130 mmHg and diastolic blood pressure ≤ 85 mmHg. Patients with T2D, impaired glucose transport, hypertension, hypercholesterolemia or fatty liver disease, as well as those undergoing drug treatments against any of these disorders, were automatically considered metabolically unhealthy (MU) regardless of their biochemical measurements. Further classification of the patients into four groups (metabolically healthy obese, MHO; metabolically healthy non-obese, MHNO; metabolically unhealthy obese, MUO and metabolically unhealthy non-obese, MUNO) was performed on the basis of obesity diagnosis (BMI ≥ 30 kg/m^2^).

### Taxonomic and functional profiling

As the curatedMetagenomicData R package provides taxonomic and functional data, we did not need to process samples from the FengQ_2015 and KarlssonFH_2013 datasets. As for the MetaHit subjects, we followed the same pipeline as in this package. Raw paired-end reads were downloaded from the European Nucleotide Archive. Reads belonging to the same sample were processed together, and two samples containing less than 10^7^ total reads were discarded. Read trimming and removal of host reads was performed with the default configuration of the *read_qc* module from metaWRAP (version 1.3.2) [31]. This pipeline uses Trim Galore to remove Illumina adapters and trim low quality ends (Phred score < 20) [32]. Next, it removes host contamination with BMTagger, for which we used the hg38 version of the human genome [33]. This tool also generates FASTQC quality reports before and after processing each sample [34].

The taxonomic composition of the microbial communities was calculated with MetaPhlAn3 [35]. The minimum read length was set to 40 nucleotides for the healthy MetaHit cohort, and default parameters were used otherwise. Relative abundance tables were agglomerated to the species level using the *tax_glom* function from the phyloseq R package (version 1.36.0) [36] and merged with those from the FengQ_2015 and KarlssonFH_2013 datasets. The resulting table was adjusted for batch effect with the *adjust_batch* function from the MMUPHin R package (version 1.6.2) [37].

Functional profiling was performed with HUMAnN3 [35] using the full UniRef90 database [38], obtaining MetaCyc pathway abundances and pathway coverages [39]. Pathway coverage files represent information as the presence (1) or absence (0) of a pathway, regardless of its relative abundance. These files were processed with MMUPHin for batch effect correction and imported to STAMP software (version 2.1.3) for visualization [40]. Differences between groups were tested using Welch’s *t*-test for two groups (MU *vs*. MH) and corrected for multiple testing via Benjamini-Hochberg FDR, with a cutoff of *q* < 10^−5^.

### Metagenomic analyses

Alpha diversity measurements reflect the diversity of species found within a sample [41]. For this analysis, we used functions from the mia R package (version 1.0.8) [42] to compute the Chao1 index for richness (function *estimateRichness*) and Shannon and Simpson’s indices for diversity (*estimateDiversity*). The Chao1 index is an indicator of the number of observed species or richness, while Shannon’s and Simpson’s indices consider species abundances as well [41]. The Shannon’s diversity index is strongly influenced by species richness and by less common species, while the Simpson’s index gives more weight to more common species [43].

We also computed Pielou’s evenness (*estimateEvenness*), the relative index for dominance (*estimateDominance*) and log-modulo skewness for rarity (*estimateDiversity*). Evenness indices focus on species abundances, and are highest when all the species have the same abundance. Pielou’s evenness is Shannon’s diversity normalized by the observed richness [41]. Dominance represents the influence that one or a few species have on the total species abundances. The relative index is a dominance metric defined as the relative abundance of the most abundant species in the community [41]. Finally, rarity characterizes the concentration of the less common taxa. Log-modulo skewness is a rarity measurement based on the skewness of the frequency distribution of relative abundances [41,43].

The Kruskal-Wallis test was used to test differences between multiple groups and the Wilcoxon rank sum test was used for pairwise tests. All *p*-values were corrected for multiple comparisons using the Benjamini-Hochberg false discovery rate (FDR) correction.

The phyloseq R package (version 1.36.0) [36] was used for beta diversity measures. These measurements reflect how different two microbial communities or samples are in their composition and structure, and are usually represented as distance matrices [41]. The *distance* function was obtained to obtain both phylogenetic (weighted and unweighted UniFrac) and non-phylogenetic (Bray-Curtis and Jaccard) measurements. The former measurements use phylogenetic information to compare the samples, while the latter do not [41]. Moreover, these metrics can either be based only on the presence or absence of each species, as is the case for unweighted UniFrac and Jaccard indices; or use the relative abundances information too, as weighted UniFrac and Jaccard indices do [41].

Dimensionality reduction of the distance matrices was performed by principal coordinate analysis (PCoA) via the *ordinate* function. Differences between groups were tested by permutational multivariate analysis of variance (PERMANOVA) with 999 random permutations using the function *adonis* from the vegan R package (version 2.5.7) [44].

### Preprocessing of relative abundances before machine learning

The relative abundance table, containing 710 features, was filtered to discard species with relative abundances lower than 0.05% in more than half of the samples. A pseudocount equal to half the minimal non-zero value was added to remove zero abundance values and a centered log-ratio (CLR) transformation was performed using the *transform* function from the microbiome R package (version 1.14.0) [45]. 117 markers with an absolute Pearson correlation coefficient greater than 0.9 were also removed. The number of metagenomic features remaining after these steps is 543.

### Random Forest

The caret R package (version 6.0.90) was used to build random forest (RF) models [46]. RFs are ensemble methods based on decision trees. Ensemble learning methods build several classifiers, and use their majority voting as the final output. In the case of RF, this is done by building each tree on a bootstrap sample of the training data [47].

Data were randomized into a training set and a testing set with a 75/25% split. Four different models were trained: two binary models and two multiclass models, trained either only on microbiome data or on microbiome data together with age, sex, and BMI information. As the BMI is used to define obesity status, and we wanted the multiclass classifier to differentiate obese from non-obese patients based only in their GM, the BMI was not included as a feature in this model. The class labels used for prediction were MH and MU for the binary model and MHO, MHNO, MUO and MUNO for the multiclass model. Models were trained by 5 repeats of 10-fold cross-validation while performing grid search for the *mtry* hyperparameter (23, 153, 283, 413, 543) and testing for different values of *ntree* (200, 600, 1000, 1400, 1800). The impurity decrease at each split was calculated via the Gini index criterion. The optimal combination of hyperparameters was chosen based on the model’s accuracy.

Then, the four tuned models were trained with different subsampling procedures: i) using the original training data, ii) using an upsampled version of the training data, and iii) using a downsampled version of the training data. These techniques try to reduce the effect of class imbalances, i.e., different numbers of samples for each class. Upsampling samples the minority class with replacement, and downsampling does the opposite with the majority class. The RF model was trained with the randomForest implementation available in the caret R package.

Finally, we predicted labels on the test data using the four chosen models, we plotted their receiver operating characteristic (ROC) curves and calculated the area under the curve (AUC) as a quality measure. This procedure was also performed with a randomized version of the class labels. For multiclass models, ROC curves were plotted for every possible class combination. ROC curves and their corresponding AUCs were calculated using the pROC package in R (version 1.18.0) [48]. Finally, feature importance was estimated with the *varImp* function from caret, which uses out-of-bag samples for prediction error estimation.

### Model validation and biomarker identification

The binary model based exclusively on metagenomic data was chosen for validation and biomarker search. The hyperparameters were fixed to *mtry* = 23 and *ntree* = 1000, and no subsampling was performed. Then, CLR-transformed relative abundances of the 20 most important features from this model were tested for significant differences between MU and MH patients via a Wilcoxon rank sum test, followed by Benjamini-Hochberg correction.

This model was also used to predict metabolic status on a cohort of 39 celiac disease patients, composed of 28 MH and 11 MH subjects (unpublished data from the host group); as well as on the 93 CRC and adenoma subjects from the FengQ_2015 dataset, which included 83 MU and 10 MH patients.

### Patient similarity network

Patient similarity networks represent patients as nodes or vertices. They are connected by an edge depending on a similarity measure of choice. We used Aitchison distances, i.e., Euclidean distances between CLR-transformed relative abundances, as a similarity measure for GM composition. The *make_network* function in phyloseq was used, setting the *max*.*dist* parameter to 65 and Aitchison distances as edge weights, and removing isolated nodes.

Network parameters such as order, size, density, diameter, average path length and clustering coefficient, as well as maximum and mean degree, were calculated with the igraph R package (version 1.2.10) [49]. Network order and size are defined as the number of vertices and edges respectively. The degree of each node represents how many nodes it is connected to. Graph density is the ratio between the number of edges in a graph and the maximum number of edges that the graph could have. The diameter of a graph is the maximum distance between any pair of vertices [50]. The average path length is the average distance between any pair of nodes. The clustering coefficient, or transitivity, for each vertex is obtained as the ratio between the number of edges that connect its neighbors and the number of edges that could exist between them. The transitivity of the whole graph is the average value of this measurement along all of its vertices [51]. We also calculated the node degree and node betweenness distribution. The betweenness of a single vertex is the fraction of the shortest paths connecting all pairs of nodes that include that particular vertex [52]. All these measurements were estimated for the patient similarity network and for an equivalent random graph, generated via the Erdos-Renyi model [53] with the function *erdos*.*renyi*.*game*.

The *assortativity* function was used to estimate the assortativity coefficient for patient data (metabolic and/or obesity status, age and study of origin), for the biomarkers obtained from the machine learning pipeline, and for alpha diversity measures. This coefficient can be interpreted as a Pearson correlation coefficient of a selected property between every pair of connected nodes [54]. To assess the robustness of these values, we obtained 100 randomized versions of the network and calculated the assortativity coefficients for each feature. Randomized graphs were built using the *rewire* function from igraph with the method *each_edge*, which reconnects each edge with a constant probability. The fraction of the randomized networks whose assortativity coefficient is equal or higher than that of the original graph for a specific feature can then be interpreted as an empirical *p*-value. These *p*-values were corrected for FDR using the Benjamini-Hochberg correction.

We then searched for communities within the network using the *cluster_fast_greedy* function, which implements a fast greedy algorithm based on edge weight [55]. Features were tested for significant differences between communities by the Wilcoxon rank sum test in the case of numerical features or using the chi-square test for categorical features, correcting all *p-*values for multiple testing via Benjamini-Hochberg FDR.

Cytoscape software (version 3.9.0) was used for network visualization [56].

### Linear Discriminant Analysis of Effect Size (LEfSe)

We used the *CRC WGS dataset*, curated by members of the Working Group 3 of the COST Action CA18131, an European network working in statistical and machine learning techniques in human microbiome studies. This dataset is composed by a collection of Metagenomic Species Pan-genomes (MSPs) obtained from 1593 patients from 10 public CRC WGS cohorts. MSPs were processed with the LEfSe Conda version 1.0.0 [57]. An MSP corresponding to *Homo sapiens* was removed from the table before running LEfSe. LEfSe was run using an alpha cutoff of 0.05 for feature significance and an effect size cutoff of 3.5.

### Data and code availability

Reads from the MetaHit project were downloaded from the European Nucleotide Archive using the accession numbers PRJEB2054 and PRJEB5224. Metagenomic, functional and patient data for the other two studies are available in curatedMetagenomicData. Additional metadata can be obtained from the online version of the articles describing each dataset [17,27–29]. The code used to analyze the data and produce our results is available in the following GitHub repository: https://github.com/blacruz17/tfm.

## Results

### Study population

Our initial search retrieved a total 7277 stool samples from 20 different studies (Table 1). Even though relevant information for patient classification had been collected according to these studies’ methods, we were restricted by the availability of such data. Most of the metagenomic data was easily accessible through the ENA, but in nearly all cases patient metadata was not public. As we needed this information to classify patients into subgroups, our population only includes 356 samples, from the studies where the relevant metadata was accessible to us. We also discarded 4 studies because, even though the information was publicly available, we wanted to avoid variation due to geographical location [58].

**Table 1.** Summary of the studies retrieved by the initial search. Datasets included in our study population are highlighted and in bold.

| Study | Method | Sample size | Country | Disease | Limitation |
| --- | --- | --- | --- | --- | --- |
| Chávez-Carbajal A [59] | 16S rRNA | 67 | Mexico | MetS <sup>1</sup> | Metadata unavailable |
| de la Cuesta-Zuluaga J [60] | 16S rRNA | 442 | Colombia | Cardiometabolic health | Metadata unavailable |
| **Feng Q [17]** | **WGS** | **156** | **Austria** | **CRC** | **Used (only controls)** |
| Jie Z [61] | WGS | 331 | China | CVD | Not European |
| **Karlsson FH [27]** | **WGS** | **145** | **Sweden** | **T2D** | **Used** |
| Kim M [62] | 16S rRNA | 747 | South Korea | MH/MU | Metadata unavailable |
| Le Chatelier E [14] | WGS | 292 | Denmark | Obesity | Metadata unavailable |
| Li J [63] | WGS | 196 | China | Hypertension | Metadata unavailable |
| Liu R [64] | WGS | 223 | China | Obesity | Metadata unavailable |
| **MetaHit Healthy [28]** | **WGS** | **85** | **Denmark** | **Healthy** | **Used** |
| **MetaHit T2D [29]** | **WGS** | **65** | **Denmark** | **T2D** | **Used** |
| Olivares P [22] | 16S rRNA | 109 | Brazil | MHO/MUO | Reads and metadata unavailable |
| Org E [65] | 16S rRNA | 531 | Finland | MetS | Metadata unavailable |
| Qin J [15] | WGS | 382 | China | T2D | Not European |
| Ross M [66] | 16S rRNA | 63 | Mexican-American | Obesity, T2D | Not European |
| Sankaranarayanan K [67] | 16S rRNA | 37 | USA | T2D | Not European |
| Zeevi D [68] | 16S rRNA | 800 | Israel | T2D | Metadata unavailable |
| Zeng Q [69] | 16S rRNA | 1914 | China | Obesity | Metadata unavailable |
| Zhong X [23] | 16S rRNA | 382 | Ireland | MetS | Metadata unavailable |
| Zupancic ML [70] | 16S rRNA | 310 | Amish (USA) | MetS | Metadata unavailable |

<sup>1</sup> Metabolic syndrome.

Thus, our samples come from 4 WGS studies, with a total of 83 MHNO, 154 MUNO, 43 MHO and 76 MUO subjects. The summary statistics for each group are shown in Table 2. There were significantly fewer women in the MUO group when compared to the MUNO category. Healthy groups were significantly younger than their unhealthy counterparts (MHNO *vs*. MUNO and MHO *vs*. MUO). These groups also showed differences in biochemical parameters, with healthy individuals having higher HDL cholesterol as well as lower fasting glucose and triglycerides. This also happened with the MUNO group compared to the MUO individuals. As for BMI, non-obese groups had generally lower values than their obese pairs, and the same happened for MHNO individuals when compared to MUNO ones. Briefly, MH subjects show lower metabolic indicators in most cases, with the exception of HDL cholesterol, as expected.

**Table 2.** Summary statistics of the study population.

| Variable | N | Lean |  | Obese |  | p-value<sup>2</sup> | q-value<sup>3</sup> |
| --- | --- | --- | --- | --- | --- | --- | --- |
|  |  | MHNO, N = 83<sup>1</sup> | MUNO, N = 154<sup>1</sup> | MHO, N = 43<sup>1</sup> | MUO, N = 76<sup>1</sup> |  |  |
| Age (years) | 356 | 64 (54, 70) | 70 (66, 71) | 59 (52, 69) | 68 (63, 70) | <0.001 | <0.001 |
| Sex | 356 |  |  |  |  | 0.045 | 0.045 |
| Female |  | 58 (70%) | 116 (75%) | 27 (63%) | 44 (58%) |  |  |
| Male |  | 25 (30%) | 38 (25%) | 16 (37%) | 32 (42%) |  |  |
| BMI (kg/m<sup>2</sup>) | 356 | 23.1 (21.7, 25.1) | 25.6 (23.0, 28.1) | 33.2 (31.7, 35.4) | 32.0 (31.0, 34.2) | <0.001 | <0.001 |
| HDL (mmol/L) | 203 | 2.07 (1.79, 2.37) | 1.66 (1.33, 2.01) | 1.69 (1.44, 1.83) | 1.37 (1.15, 1.71) | <0.001 | <0.001 |
| Fasting glucose (mmol/L) | 205 | 5.30 (4.95, 5.70) | 5.83 (5.40, 6.70) | 5.50 (5.30, 5.80) | 6.47 (5.82, 7.43) | <0.001 | <0.001 |
| Triglycerides (mmol/L) | 204 | 0.91 (0.71, 1.16) | 1.07 (0.88, 1.56) | 1.01 (0.69, 1.15) | 1.55 (1.20, 2.09) | <0.001 | <0.001 |

<sup>1</sup> Median (IQR); n (%)
<sup>2</sup> Kruskal-Wallis rank sum test; Pearson's Chi-squared test
<sup>3</sup> False discovery rate correction for multiple testing

All datasets were processed through the same pipeline: quality trimming and removal of host reads followed by taxonomic profiling with MetaPhlAn3 and functional profiling with HUMAnN3. As the study of origin showed a significant effect on the variance between patients, species relative abundances were corrected for batch effect (Figure 1). This reduced the variability explained by the study from 12.73 % to 2.60 %, although its effect is still significant (*p* < 0.001, PERMANOVA). The same process was performed on pathway files (Supplementary Figure 1).

**Figure 1.**
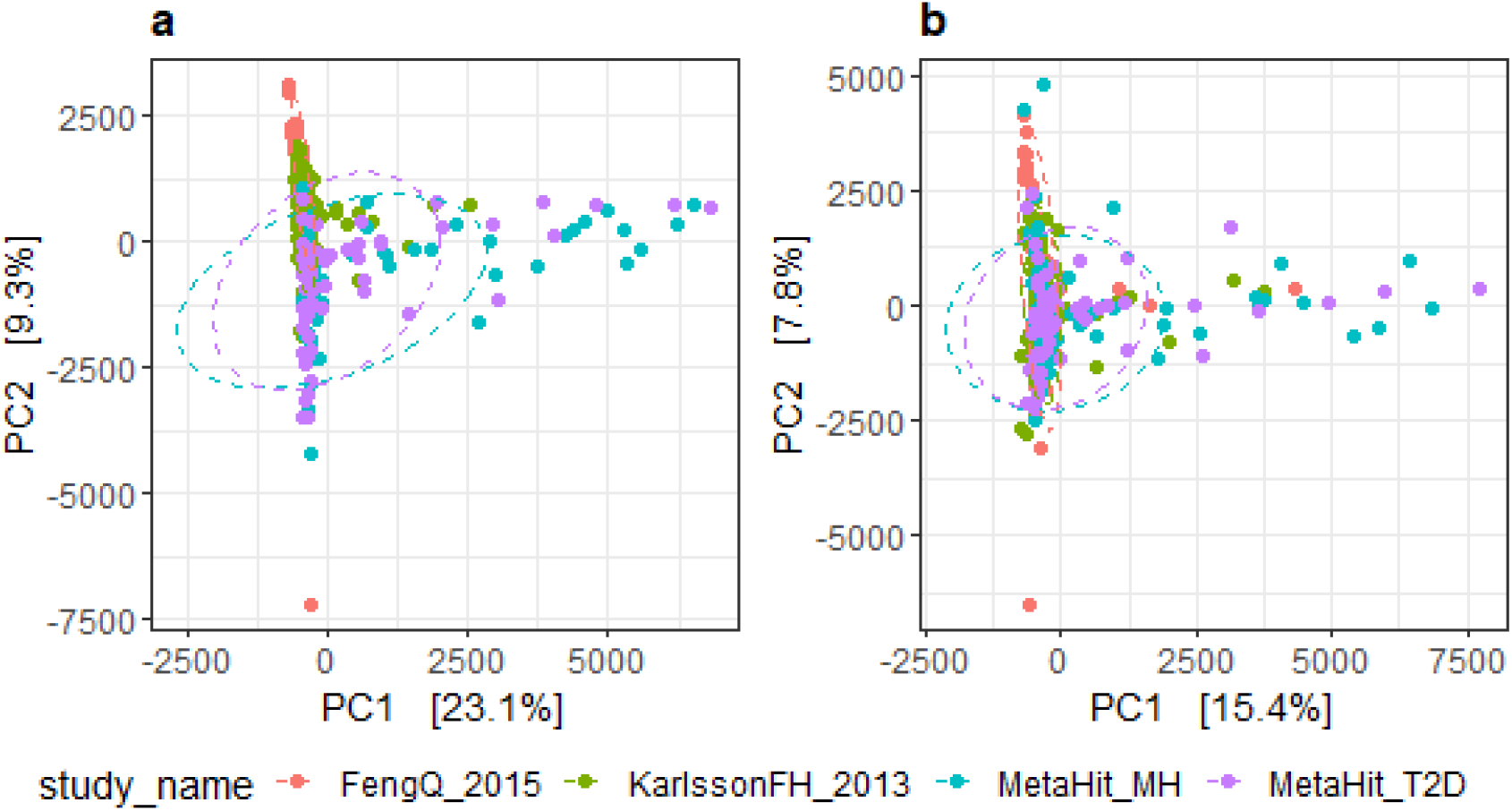
Batch effect correction with MMUPHin. Samples were ordinated via principal component analysis (PCA) and the two first components were plotted before (**a**) and after (**b**) batch effect correction with MMUPHin. Ellipses represent 95% confidence intervals for each group.

### Community diversity: alpha diversity

We did not find any significant differences in community diversity when testing samples from each study separately. When all samples were analyzed together, we found that the MUNO group showed significantly higher diversity than the MHO group as measured by the Shannon and Simpson indices, while dominance and rarity values were significantly lower in the MUNO group. Additionally, MHNO subjects showed significantly higher dominance than the MUNO group (Figure 2). We also compared patients only by obesity or by metabolic status (Supplementary Figure 2). No measures were significantly different between obese and lean patients, while the MU group showed higher values for Shannon and Simpson’s indices and a lower relative index than the MH group. Summing up, we found that MU patients generally had more diverse gut microbiomes than their healthy pairs.

**Figure 2.**
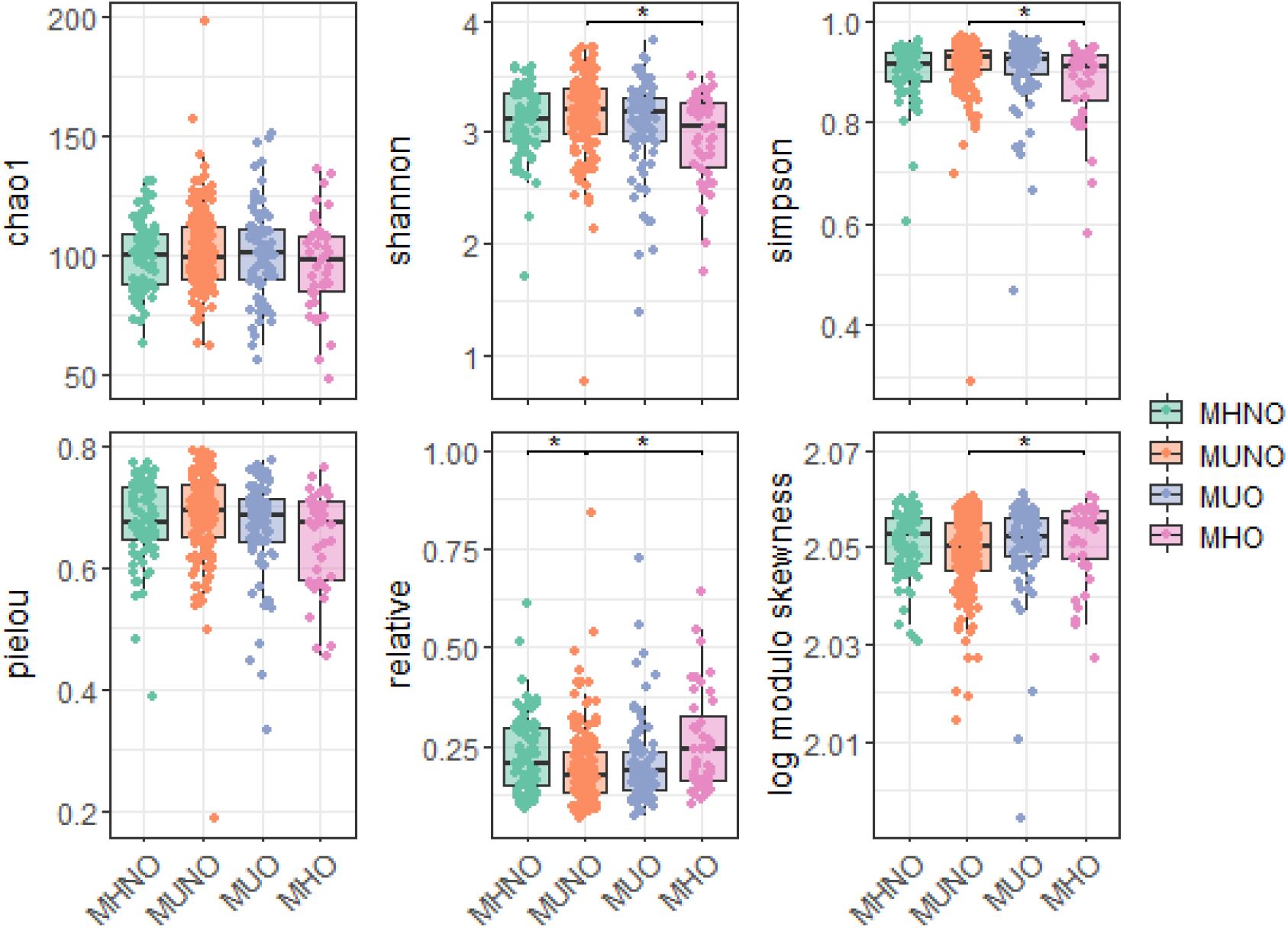
Alpha diversity differences between groups. The Shannon and Simpson diversity indices showed significant differences in the Kruskal-Wallis rank sum test (*p* = 0.0417 and *p* = 0.0134 respectively), as well as the relative index (*p* = 0.0134) and the log-modulo skewness (*p* = 0.0417). These same measures were significantly different between MUNO and MHO groups (*p* = 0.024, *p* = 0.024, *p* = 0.024 and *p* = 0.0432 respectively). MHNO and MUNO groups showed differences in the case of the relative index (*p* = 0.042).

### Differences between communities: beta diversity

In order to evaluate differences in community structure and composition between samples, we calculated both phylogenetic (UniFrac weighted and unweighted distances) and non-phylogenetic (Bray-Curtis, Jaccard) indices. Even though we found significant differences between groups as assessed by PERMANOVA (*p* < 0.001), their explained variance was very low, ranging from 1.60 % to 3.89 %. As a result, groups could not be clearly separated by PCoA, as can be seen in Figure 3.

**Figure 3.**
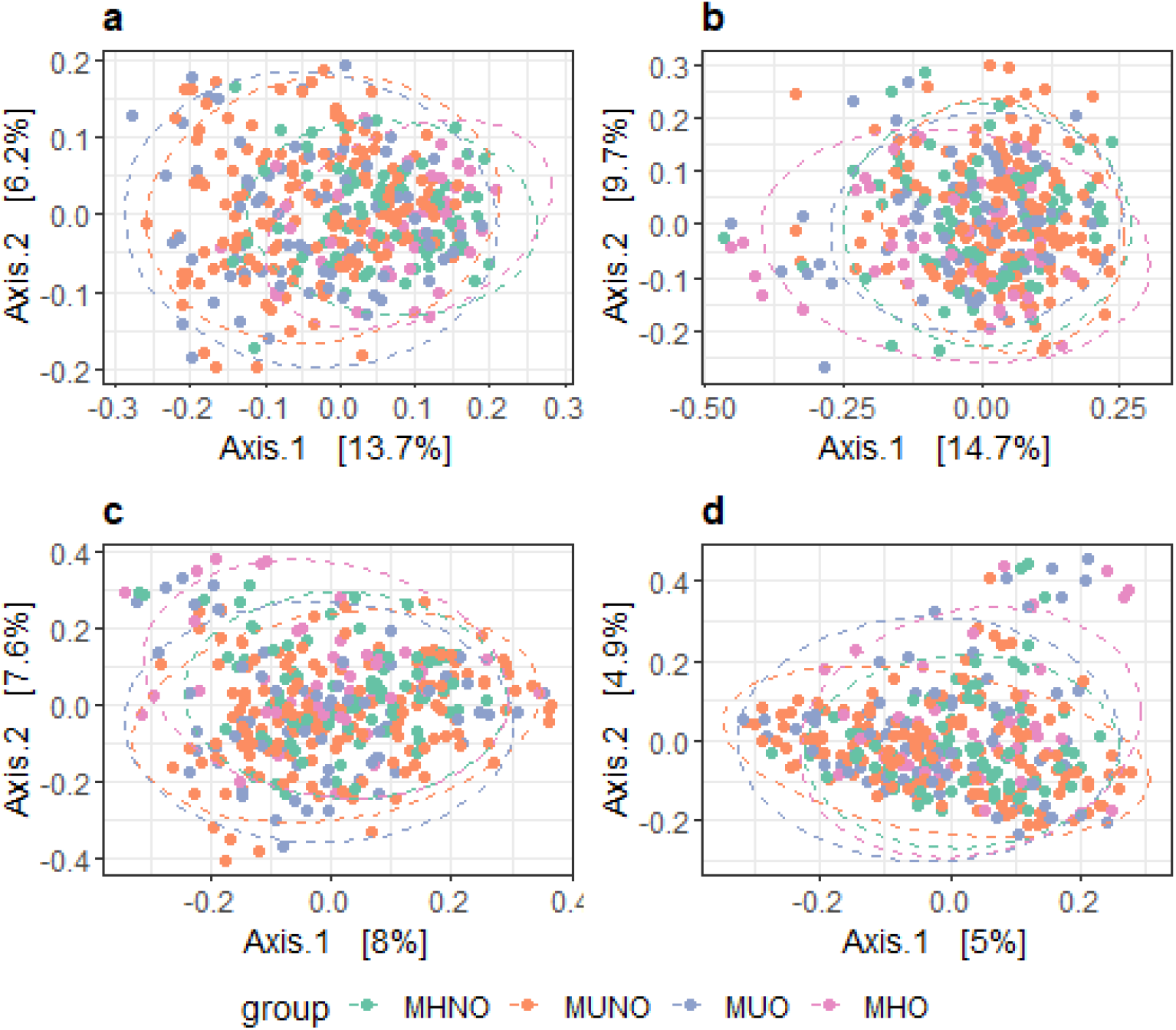
Beta diversity analysis: PCoA plots. Phylogenetic (**a-b**) and non-phylogenetic (**c-d**) distance measurements were calculated with phyloseq and represented via PCoA. **a)** unweighted UniFrac, **b)** weighted UniFrac, **c**) Bray-Curtis, **d**) Jaccard index. Differences between groups were tested with PERMANOVA with 999 permutations. Ellipses represent 95% confidence intervals.

### Functional profiling

We used STAMP to visualize functional differences between MH and MU patients. There were two pathways with significantly different coverage and with an effect size greater than 1: the superpathway of fatty acid biosynthesis initiation in *Escherichia coli* (FASYN-INITIAL-PWY) and the peptidoglycan maturation (meso-diaminopimelate containing) pathway (PWY0-1586), both shown in Figure 4.

**Figure 4.**
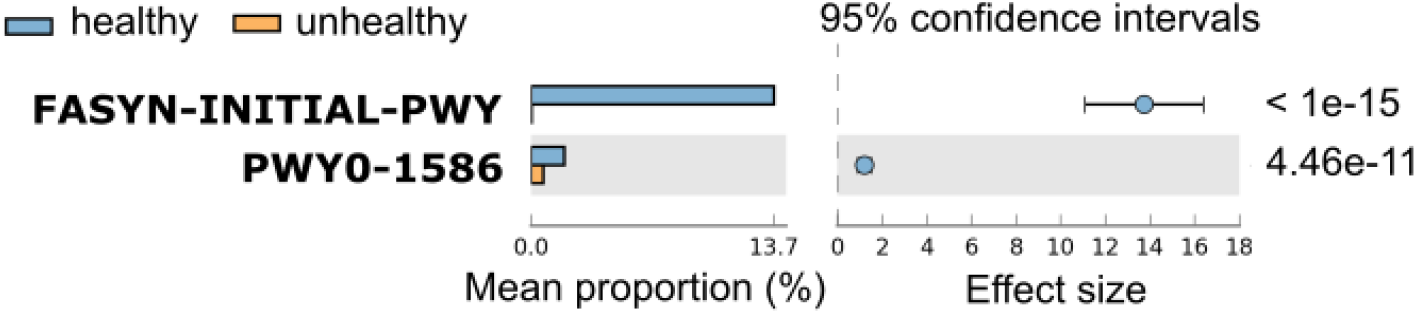
STAMP visualization after functional profiling with HUMAnN3: pathway coverages. Extended error bar plot of features showing a corrected *p*-value < 10^−5^ and an effect size > 1. Mean proportions are shown as percentages. Effect size is measured as the difference between proportions. FDR-corrected *p-*values for each pathway are shown on the right.

### Random Forest classifier based on gut microbiota composition

RF classifiers are widely used in microbiome studies [71,72]. Previous studies have found them to perform better than other classical machine learning algorithms such as support vector machines or regularized logistic regression [73]. Furthermore, they perform better than deep learning approaches in datasets with small sample sizes [71]. Finally, they can support multiclass classification tasks natively [72], making them a suitable solution for our problem. We built a multiclass RF classifier to further investigate the potential of the GM to classify MHO and MUO patients, using the CLR-transformed relative abundances table as input data. The low proportion of MHO subjects in the dataset led us to construct a binary model as well, with the purpose of discriminating metabolically healthy *vs*. unhealthy subjects irrespective of the subject’s obesity status.

We trained the models by 5-times 10-fold cross-validation. The accuracy reached during the cross-validation process and the AUC over the test set were used as measures of model quality. We tested different values for two hyperparameters: the number of features that are considered at each split (*mtry*) and the number of trees that are built for the ensemble (*ntree*) (Figure 5). The combination reaching the highest accuracy during cross-validation was chosen and the resulting models were used to predict class labels on the test set. Resultsm for the models trained on a dataset including the patient’s age, sex, and BMI as well as their metagenomic taxonomic profiles are shown in Supplementary Figures 3 and 4.

**Figure 5.**
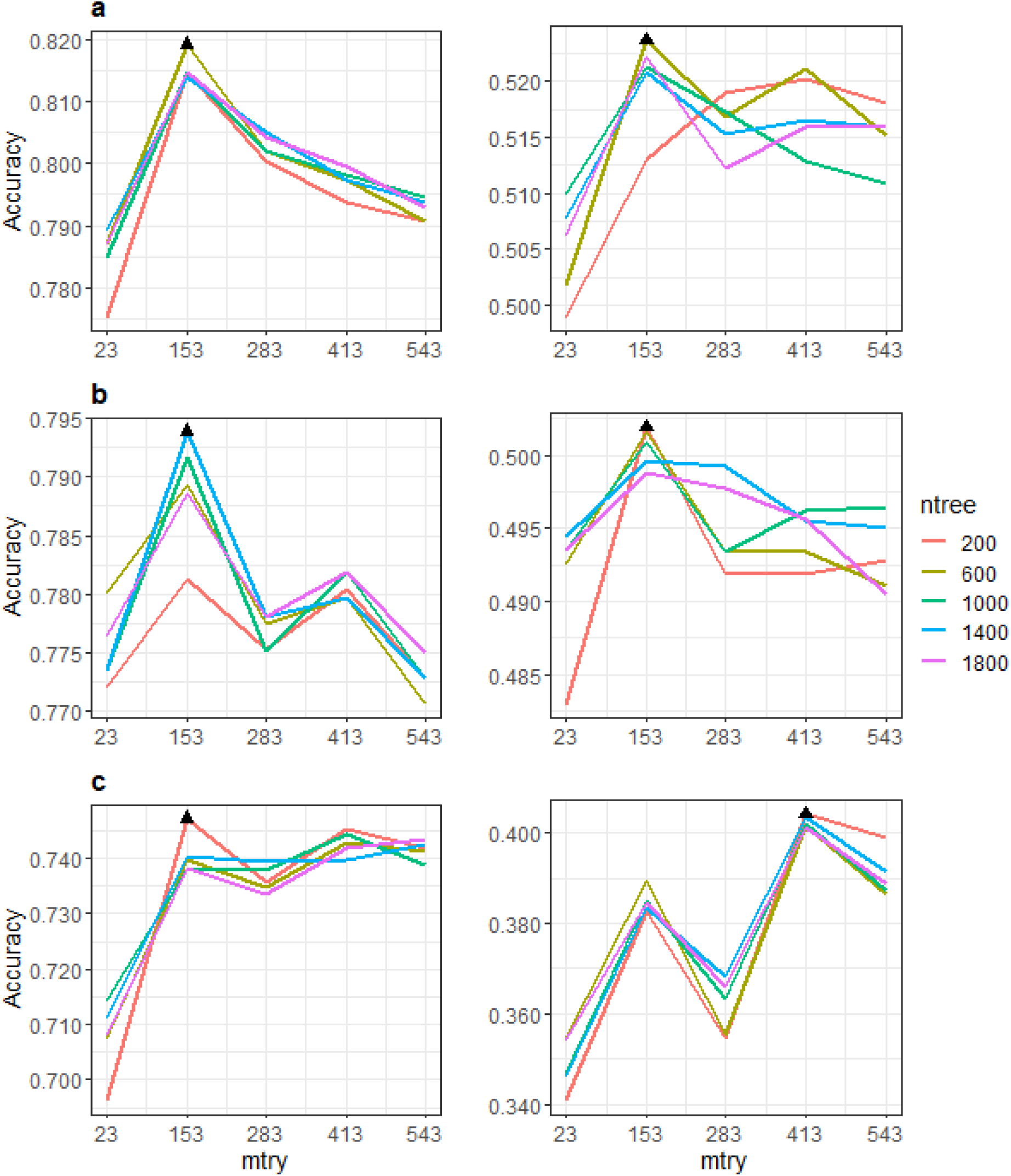
Hyperparameter tuning for abundances-based models. **a)** Models without subsampling, **b)** upsampled models, **c)** downsampled models. *mtry* and *ntree* hyperparameters were tuned in binary (left) and multiclass (right) models. Tested *mtry* values are shown in the x-axis, and different *ntree* values are represented by different line colors. Black triangles show the hyperparameter combination reaching the highest accuracy in each case.

The binary classifiers that we built to separate MU and MH patients achieved a high AUC without the need to use subsampling techniques (Fig. 6a and Supp. Fig. 4a). Thus, they successfully classified patients according to their metabolic health status, independently of their obesity status. Then, we plotted ROC curves for every class label combination in the multiclass models (Fig. 6b and Supp. Fig. 4b). These reached high AUCs for the MHO/MUO and MHO/MUNO classes (Fig. 6b and Supp. Fig. 4b) when trained exclusively on GM data. Afterwards, we trained these models including age and sex as input features as well. This maintained the high AUCs for the binary models (Supp. Fig. 4a) and for MHO/MUO and MHO/MUNO prediction in the multiclass model (Supp. Fig. 4b). Moreover, it increased the AUCs for discrimination of MHNO subjects from MUNO and MUO patients (Supp. Fig. 4b). Variable importance plots (Supplementary Figure 5) show that age became the second most important feature for patient classification in the multiclass model. In conclusion, our models allowed us to classify MU and MH patients, requiring either GM data alone or GM data together with patients’ age.

**Figure 6.**
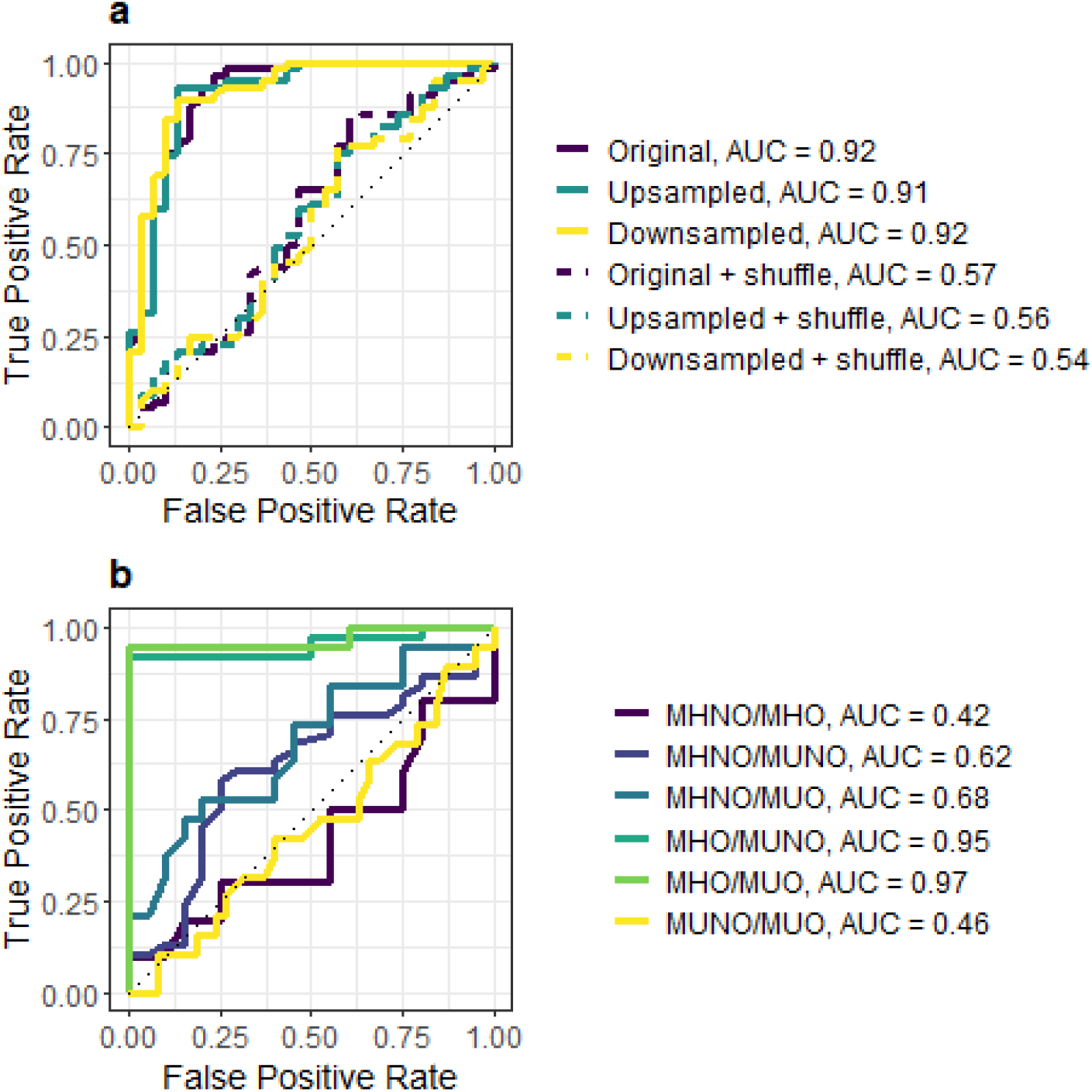
ROC curves for abundances-based models. **a)** The binary model with the best configuration for each subsampling technique, as shown in Figure 5, was used to predict class labels on the test set. Dashed lines show the ROC curves for a randomized version of the class labels. **b)** ROC curves for every possible class combination are shown for the multiclass model with *ntree* = 600, *mtry* = 153 and without subsampling.

Therefore, we used a binary RF classifier based only on CLR-transformed microbiome data for validation, with 1000 trees and an *mtry* value equal to the square root of the number of original features. As subsampling techniques did not improve the AUC, we decided not to apply any in this model. Predictions on the test yielded an AUC of 0.88 (Figure 7a). As for the validation cohorts, all subjects in a celiac disease dataset were predicted to be metabolically unhealthy, when there were actually 28 MH and 11 MU patients (AUC = 0.64). This happened with the CRC and adenoma patients from the FengQ_2015 study as well (10 MH and 83 MU patients, AUC = 0.57) (Fig. 7b).

**Figure 7.**
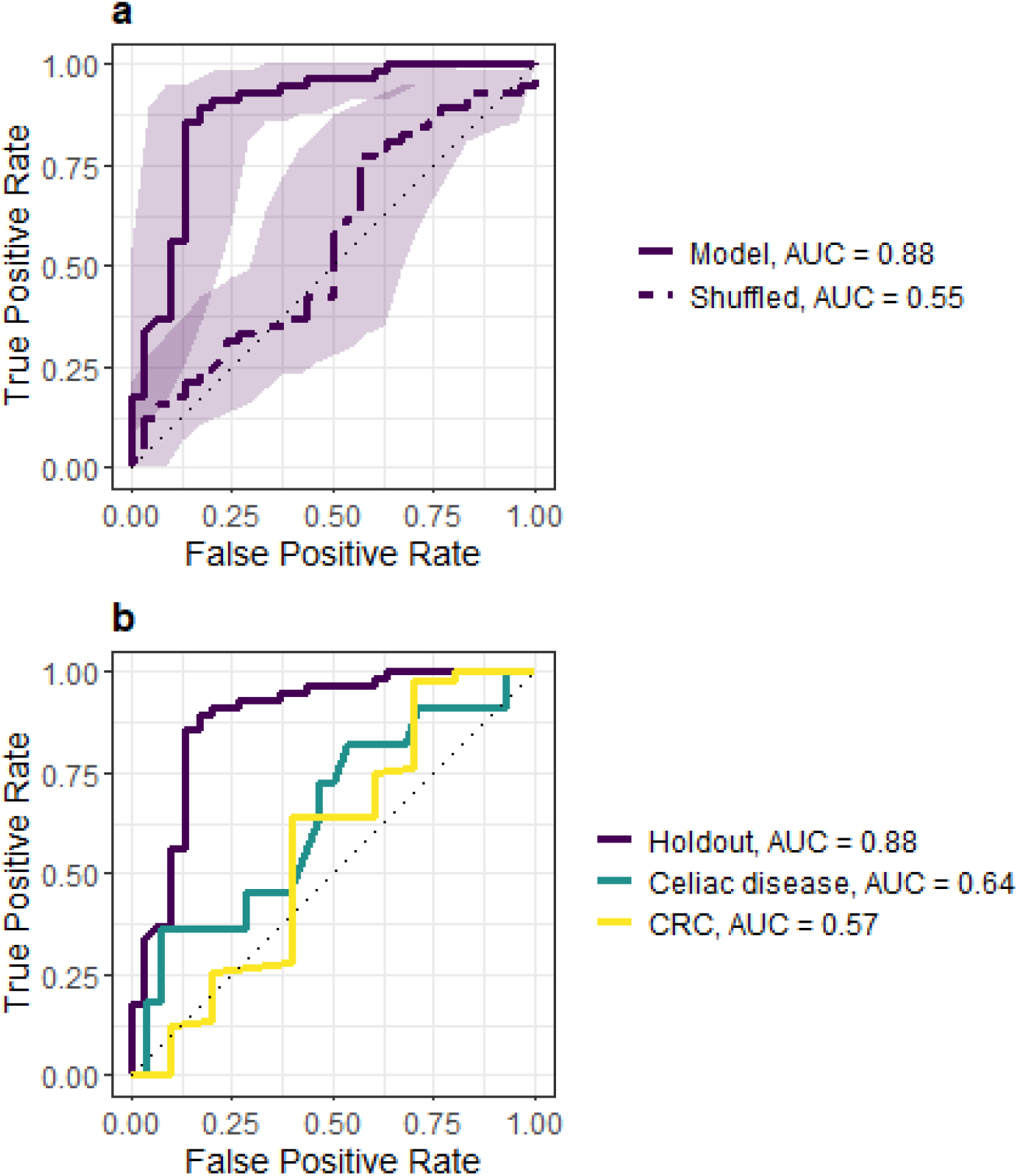
ROC curves for the final RF model. A RF model with *mtry* = 23 and *ntree* = 1000 was trained on the relative abundances table without any subsampling. **a)** ROC curves for the predictions on the holdout dataset as well as a shuffled version of the class labels. Shadowed areas represent 95% confidence intervals for sensitivity. **b)** ROC curve for the holdout dataset compared with the ROC curves for the two validation cohorts.

Biomarker search based on the final RF classifier was performed by choosing the 20 most important features in the trained model and testing them for differences in CLR-transformed abundances between MH and MU patients. Four species were detected as markers in MU patients: *Clostridium leptum* (*p* = 1.29 × 10^−4^), *Gordonibacter pamelaeae* (*p* = 4.58 × 10^−6^), *Eggerthella lenta* (*p* = 8.35 × 10^−6^) and *Collinsella intestinalis* (*p* = 5.00 × 10^−3^), sorted by descending importance in the classifier (Figure 8).

**Figure 8.**
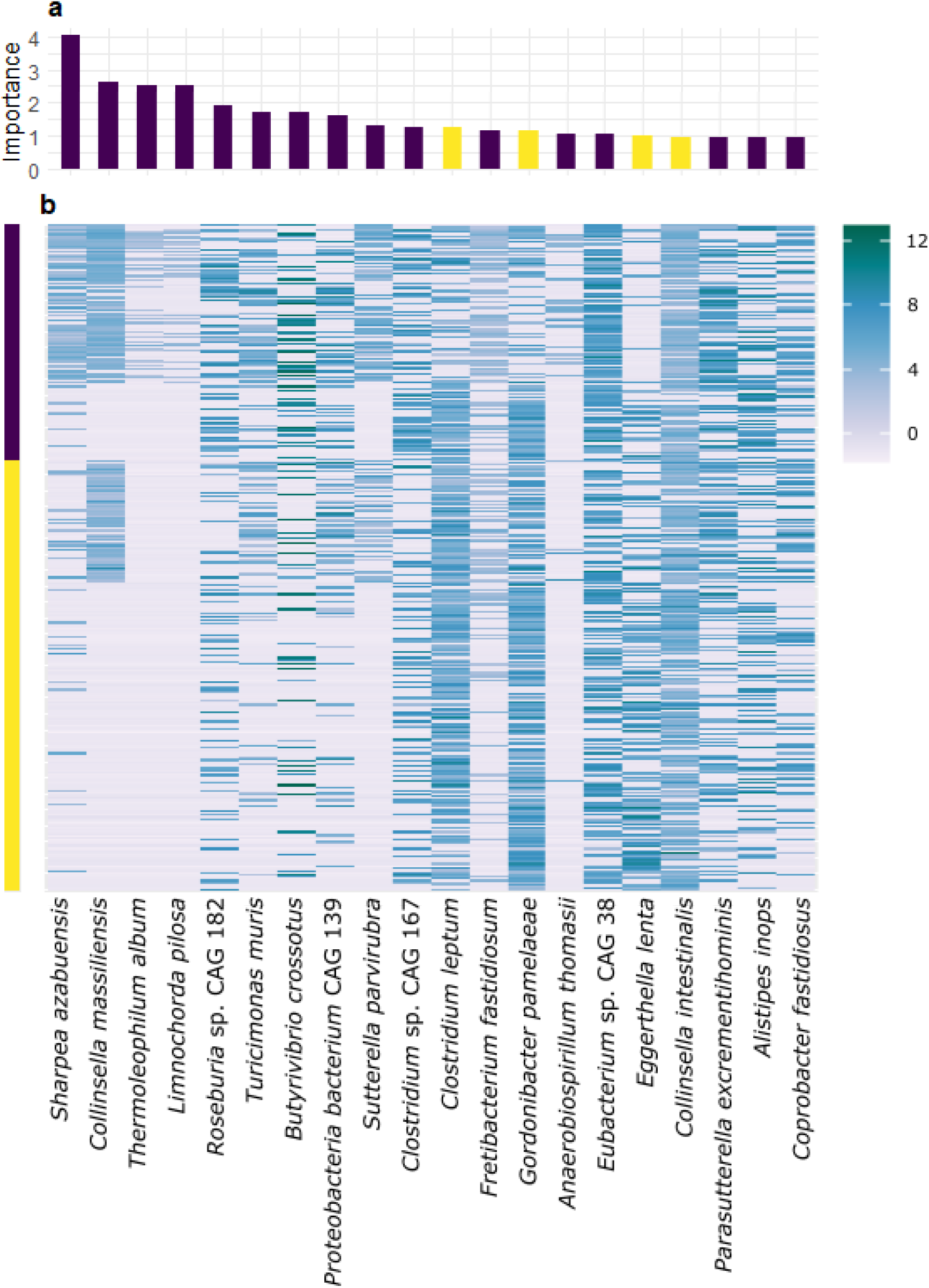
20 most important features in the final model. **a)** Feature importances. Bar height indicates feature importance and bar color indicates whether the marker is enriched in MH (purple) or MU (yellow) patients. **b)** Heatmap of CLR-transformed relative abundances. The left bar indicates the patients’ metabolic category. Purple: MH, yellow: MU.

### Patient similarity network

Patient similarity networks have arisen as an alternative to machine learning methods for patient classification. Their main advantage is that they are more easily interpreted than machine learning algorithms, which is a valued quality in clinical models [74]. In order to assemble a patient similarity network, we calculated Aitchison distances between patients. After trying different cutoff values (Table 3), we set the maximum distance between patients to 65, so that only patients with an Aitchison distance ≤ 65 between them were connected by an edge. As a consequence, only 53 isolated nodes were removed, allowing us to maintain 303 patients in the network. These represent 85 % of the samples, thus keeping most of the information. The resulting network has 5457 edges, out of the 63190 edges that connect all patients with each other.

**Table 3.** Network size, order, number of communities and order of the largest community using different cutoff values for the maximum Aitchison distance between nodes.

|  | Whole | 80 | 75 | 70 | 65 | 60 | 55 | 50 |
| --- | --- | --- | --- | --- | --- | --- | --- | --- |
| **Size** | 63190 | 47979 | 33823 | 17406 | 5457 | 886 | 64 | 3 |
| **Order** | 356 | 356 | 356 | 348 | 303 | 214 | 61 | 6 |
| **No. of communities** | 2 | 2 | 2 | 3 | 5 | 9 | 13 | 3 |
| **Largest community** | 355 | 195 | 184 | 171 | 151 | 68 | 9 | 2 |

Our network consists of a single connected component, with an average of 36 neighbors for each node, as illustrated in Table 4 together with other network properties. We also evaluated node betweenness and node degree distribution for this graph (Figure 9) and for an equivalent random graph. The clustering coefficient and average path length of our network are greater than those of the random graph, and there are also differences in the node betweenness and degree distributions (Table 4 and Supplementary Figure 6).

**Table 4.** Network metrics. Metrics for the patient similarity network (Network), a random equivalent graph (Random) and the five communities are shown.

|  | Network | Random | Communities |  |  |  |  |
| --- | --- | --- | --- | --- | --- | --- | --- |
|  |  |  | 1 | 2 | 3 | 4 | 5 |
| **Order** | 303 | 303 | 151 | 130 | 7 | 7 | 8 |
| **Size** | 5,457 | 5,457 | 1,697 | 1,882 | 9 | 10 | 10 |
| **Max degree** | 166 | 166 | 91 | 104 | 5 | 5 | 5 |
| **Mean degree** | 36.02 | 36.02 | 22.48 | 28.95 | 2.57 | 2.86 | 2.50 |
| **Graph density** | 0.12 | 0.12 | 0.15 | 0.22 | 0.43 | 0.48 | 0.36 |
| **Diameter** | 7 | 4 | 7 | 5 | 4 | 4 | 4 |
| **Average path length** | 2.23 | 1.89 | 2.20 | 1.86 | 1.67 | 1.57 | 1.86 |
| **Clustering coefficient** | 0.43 | 0.12 | 0.45 | 0.51 | 0.43 | 0.50 | 0.16 |

**Figure 9.**
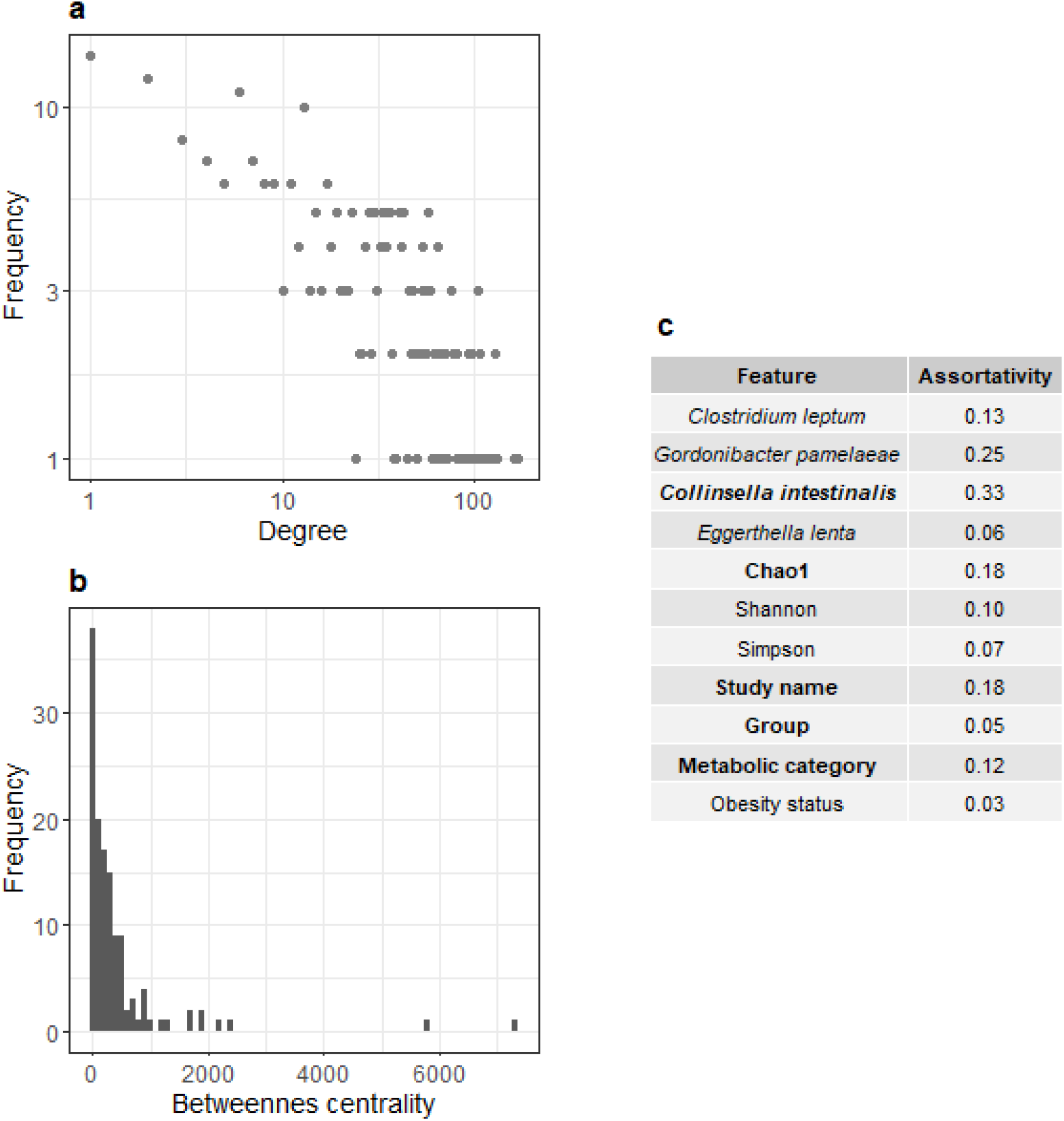
Network exploratory analysis. **a)** Scatterplot of node degree distribution. Both axes are shown on a logarithmic scale. **b)** Histogram of node betweenness centrality distribution. **c)** Assortativity coefficients for each feature. Features with an adjusted *p-*value < 0.05 are in bold.

We were also interested in whether patients with similar properties tended to interact with each other. This can be measured by the assortativity coefficient, a metric used in social sciences to measure the tendency of similar people to be connected with each other [54,75]. As an example, *C. leptum* would have a high assortativity if patients with high relative abundances were connected between them. We calculated this measure for different features, including our biomarkers of interest, several alpha diversity measures, and the patients’ metabolic or obesity status. Then, we assessed its robustness by 100 random rewirings of the network, obtaining empirical *p*-values for each feature (see Fig. 9c).

We then looked for non-overlapping communities within the network using a fast greedy algorithm. This returned a total of five communities, although the majority of the nodes (281) belonged to only two of them. Differential analysis between these two revealed that they had a significantly different number of MU and MH patients (*p* = 5.94 × 10^−7^). The community enriched in MU patients showed greater CLR-transformed abundances for three out of the four biomarkers obtained from the RF pipeline (with adjusted *p*-values of 1.34 × 10^−6^ for *C. leptum*, 1.10 × 10^−20^ for *G. pamelaeae* and 2.73 × 10^−25^ for *E. lenta*). The only exception was *C. intestinalis* (*p* = 0.156) (Figure 10). Network properties for these clusters are also shown in Table 4.

**Figure 10.**
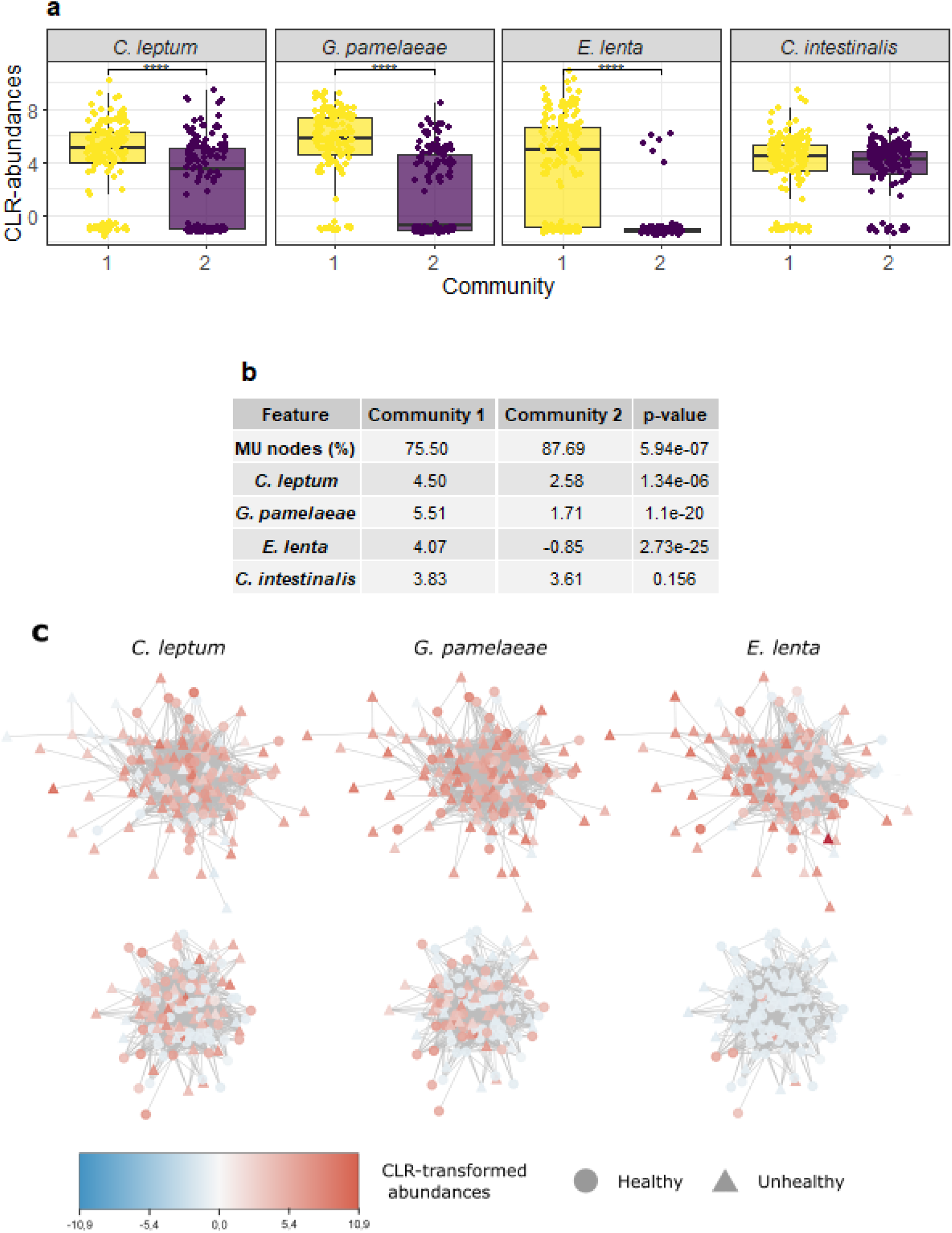
Differential analysis of the two main network communities. **a)** Boxplots showing CLR-transformed abundances for each of the MU biomarkers in each community. **b)** Table showing the proportion of MU nodes and the mean of CLR-transformed biomarker abundances for each community, together with their adjusted *p*-values (Benjamini-Hochberg). **c)** Visualization of the two main communities (top: community 1, bottom: community 2). Node shape represents metabolic health (circle: MH, triangle: MU) and node color represents CLR-transformed abundances for *C. leptum, G. pamelaeae* and *E. lenta*.

### Metabolic disease markers in a CRC cohort

Last, we evaluated whether biomarkers characteristic of MU subjects could be found in a cohort of CRC and adenoma patients. For this purpose, we performed a LEfSe analysis on a collection of 1593 fecal metagenomes collected and analyzed by the ML4Microbiome COST Action. This cohort contains 183 adenoma patients, 662 CRC patients and 748 controls. LEfSe first compares the relative abundances of all species across the different groups with a Kruskal-Wallis rank sum test. Then, significant features are used as input for linear discriminant analysis (LDA), obtaining a measure of effect size.

This analysis yielded a total of 43 features of interest, which are displayed in Figure 11 and Supplementary Figure 7. *G. pamelaeae, E. lenta* and *C. intestinalis* all belong to the family *Coriobacteriaceae*, which was detected as a marker for adenoma patients in this analysis, as well as their order (*Coriobacteriales*) and class (*Coriobacteriia*). *C. leptum* belongs to the *Clostridia* class, which is also enriched in adenoma patients.

**Figure 11.**
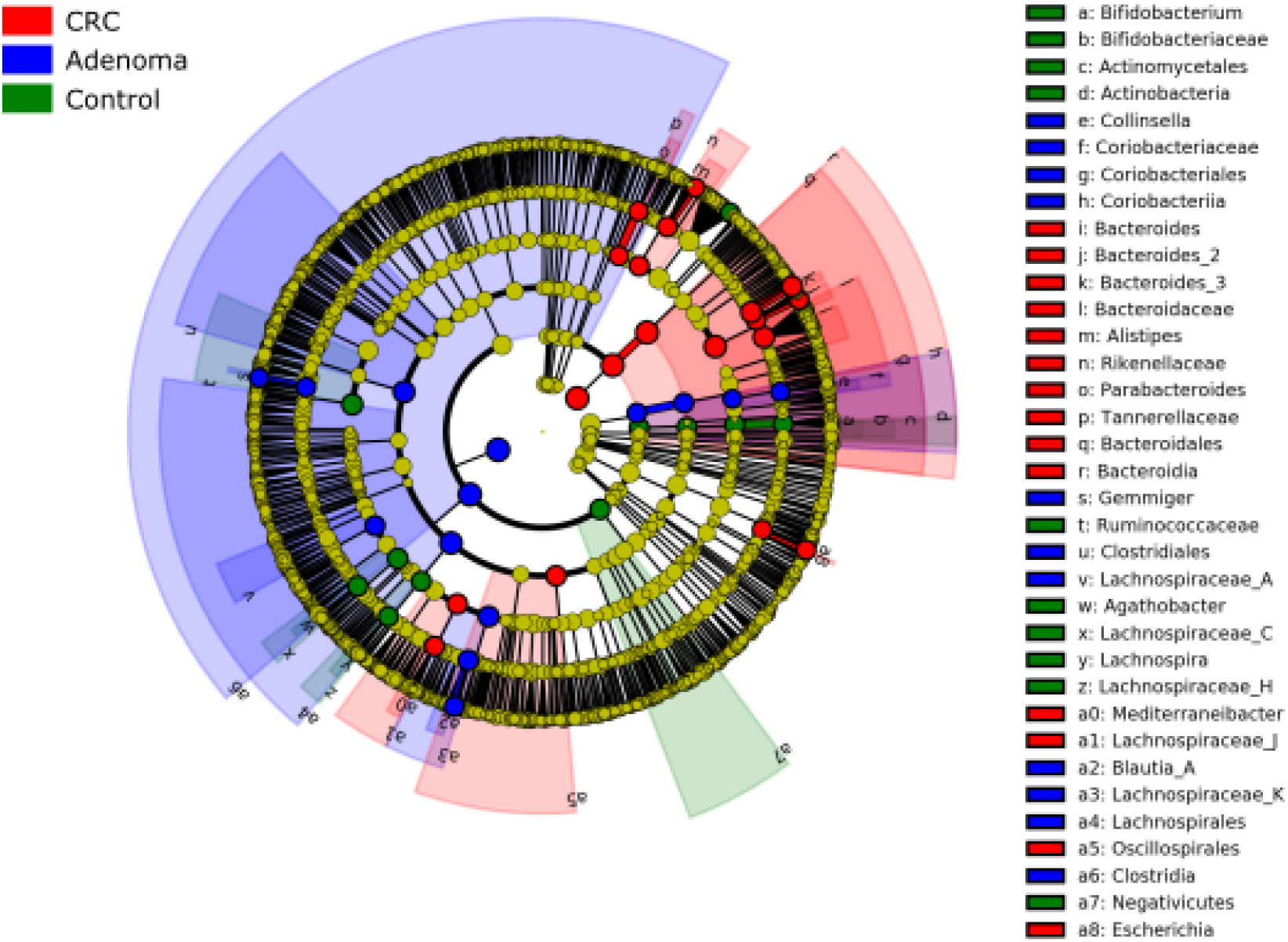
LEfSe of the ML4Microbiome COST Action patients. A dataset containing 183 adenoma patients, 662 CRC patients and 748 controls was analyzed with LEfSe. The cladogram represents the 43 significant features (*p* < 0.05, LDA > 3.5) retrieved by the analysis together with their phylogenetic relationships.

## Discussion

In this study, we analyzed the GM of 356 European patients and used it to classify them according to their metabolic health status. Individuals in the MU and MH groups had significant differences in their anthropometric parameters. These differences are due to the MU/MH classification criteria, which introduces a bias so that MU patients have increased metabolic indicators in general, and are consistent with the findings of previous studies [22,76,77]. We also found the MHNO individuals to be younger than their unhealthy pairs, consistently with the fact that the prevalence of metabolic disease increases with age [78].

Our alpha diversity analyses only found differences between the MUNO and MHO groups, making it difficult to understand the individual influences of metabolic health and obesity status. Our findings contradict previous studies that have found MUO patients to show lower alpha diversity indices than their healthy counterparts [22,23,62]. However, there are also studies which have found obese or CRC patients to have more diverse gut microbiomes than their respective controls [18,59]. Recent meta-analyses motivated by these inconsistencies in the literature state that these measurements are not reliable indicators of health or disease [79,80]. We also found differences between groups in beta diversity indices, but due to their small effect size we did not believe they were biologically relevant.

We then built a RF classifier with the purpose of classifying patients into different metabolic categories based on their gut microbiome. Multiclass classifiers separated MHO *vs*. MUO subjects successfully. Remarkably, including age as an input feature in these models improved the AUC for the classification of MUNO/MHNO patients. However, we were not able to discern obese from lean patients. Although this might be a limitation derived from our small sample size, it should be noted that there is controversy over whether metagenomic data is enough to separate obese and lean patients [79]. Some authors argue that additional information, such as metabolomic or host genomic data, might be necessary [79,81].

As, in practice, both the binary and the multiclass classifiers provided us with the same information (the metabolic status of the patients), we decided to use the binary classifier for validation, as it is simpler and its output is easier to interpret. This model classified MU and MH subjects in the holdout dataset with an AUC of 0.88. Subsequent validation of this model on two additional cohorts revealed a notable decrease in the AUC, as all patients were classified as metabolically unhealthy. This could be indicating a low generalizability of the model. However, it could also be the case that the model is actually detecting a dysfunctional GM rather than a signature specific for metabolically unhealthy patients. All patients in the validation datasets suffered from either CRC, adenoma or celiac disease, meaning that their GM is already altered. The lack of a validation cohort formed by individuals whose GM has not been reshaped by other conditions is a limitation of this study.

Nevertheless, the difficulty of identifying microbial biomarkers specific to a certain condition is not unique to this work. Previous meta-analyses that have collected case-control studies for several diseases, constructed machine learning models and performed cross-study validation have revealed that many markers associated with disease are common to more than one condition [73,82]. Moreover, some authors argue that the shift from a healthy gut microbiome towards an unhealthy one is mostly based on stochastic changes [83]. This stochasticity would lead to unstable GM states, causing dysbiotic GMs to be more diverse than healthy ones [80,83]. This phenomenon has been called the “Anna Karenina principle”, imitating the quote from Leo Tolstoy’s book: “all happy families are alike; each unhappy family is unhappy in its own way” [80,83]. Although this hypothesis is compatible with some deterministic changes [80], it could explain why it can be challenging to find biomarkers specific to a particular disease. However, there is some controversy around this matter, as it seems that this instability in the GM composition is not universal to all diseases [84].

The main challenge of using machine learning techniques in precision medicine is their interpretability, as it is important for physicians to understand how the diagnosis was made. Patient similarity networks partially overcome this limitation by providing a visualization of the decision boundary between classes [74], and have been used successfully in tasks such as the identification of cancer and type 2 diabetes subtypes [85,86]. To explore this approach, we have assembled a patient similarity network based on Aitchison distances. When we looked for communities within the graph, we found that most patients belong to two subgroups. One of them is characterized by a higher proportion of MU patients and greater abundances of the species *C. leptum, G. pamelaeae* and *E. lenta*.

The differences between MU and MH patients might be caused by the loss of microbial species associated with the MH phenotype. Six of these markers belong to the phylum *Firmicutes* (*Sharpea azabuensis, Limnochorda pilosa, Roseburia* sp. CAG 182, *Butyrivibrio crossotus, Clostridium* sp. CAG 167 *and Eubacterium* sp. CAG 38), which is enriched in short-chain fatty acid producers. These metabolites have beneficial effects on glucose homeostasis, the gut barrier and the host immune system [87]. This would support the possible role of these biomarkers in maintaining the metabolically healthy phenotype.

Alternatively, progression towards the MU phenotype may be due to an increase in certain species. Among the MU biomarkers obtained from the RF model, *C. leptum* was the most important one for patient classification. A previous study also found it to be among the top features used in a RF classifier for CRC detection [18]. Moreover, *C. leptum* has also been linked with esophageal cancer [88], and both the order *Clostridiales* and the class *Clostridia* were enriched in adenoma patients from the CRC cohort.

Our other three MU biomarkers (*G. pamelaeae, E. lenta* and *C. intestinalis*) belong to the family *Coriobacteriaceae-Eggerthellaceae* in phylum *Actinobacteria. Collinsella* spp. were enriched in tumor samples in colorectal adenoma [89], and both *Collinsella* and *Eggerthella* have been found in feces from CRC patients [19,90]. Accordingly, the family *Coriobacteriaceae*, together with the order *Coriobacteriales* and the class *Coriobacteriia*, were identified as biomarkers for adenoma in the COST Action cohort. Thus, all of the MU-related species were enriched in this group, which highlights their potential as CRC risk and progression biomarkers.

*Coriobacteriaceae* members are involved in bile acid metabolism, whose alteration has been linked not only to metabolic or gut barrier dysfunction but also to colon cancer and chronic intestinal inflammation [91–94]. There is evidence that suggests a link between a high-fat diet, an altered bile acid composition, GM dysbiosis and colitis in mice, suggesting a bridge between dietary habits, the GM and inflammatory bowel diseases [93]. *E. lenta* has also been described as an imidazole propionate producer [95]. This metabolite impairs insulin signaling and glucose tolerance and is increased in the serum of T2D patients [95]. Indeed, the prevalence of *E. lenta* has been linked with T2D [15,95].

Furthermore, a distinctive characteristic of this family is the conversion of food polyphenols such as urolithins [91,96], metabolites with a wide range of biological properties including anti-inflammatory and antioxidant effects [97]. The production of urolithins has great interindividual variability, and three urolithin metabotypes (UM) have been described based on their production. Members of the *Coriobacteriaceae* are enriched in patients belonging to the UM-B metabotype. Previous studies have identified a relationship between high cholesterol levels and this metabotype, suggesting a relationship between blood cholesterol levels, increased CVD risk and the abundance of microorganisms belonging to this family [96,98,99]. The metabolic potential of this family and its associations with disease underscore its interest in metabolic disorders research.

The main limitation of this study is the small sample size. Depending on publicly available datasets that also have the appropriate information to stratify patients according to their metabolic health and obesity status restricted the number of samples that we could use. We only included studies performed on European patients and with similar sample collection and sequencing protocols, which restricted our search even further. This strains the generalizability of our classifier. Including a greater variety of samples in our classifier might not only improve its generalizability, but also allow it to differentiate between lean and obese subjects if such a thing is possible. Another limitation derived from relying on public data is that this is a cross-sectional study, meaning that we only have information about GM composition at a single time point. Tracing its changes over time would help us better assess CRC risk, rather than depending on comparing differentially abundant species among different cohorts, as we did in our analysis.

We believe that it would be interesting to assess the robustness of our markers in future work. Some strategies that can be performed on the available data include implementing feature selection techniques in our RF model; assembling more classifiers, such as support vector machines or logistic regression; or accompanying our pipeline with additional differential analysis tools for microbiome data. The idea is that robust biomarkers would appear in more than one of these analyses. However, it would be ideal to access information from longitudinal studies that collect samples at several time points. This would allow us to confirm whether our biomarkers have a role in CRC and MUO pathophysiology, or if they are involved in general dysbiosis. Laboratory work is also necessary to validate our findings.

Investigating the MHO phenotype may improve our understanding of how obesity is associated with a greater risk of different comorbidities, especially CRC. The GM could have a decisive role in this progression. To evaluate this possibility, we have developed a RF model and a patient similarity network to separate MU and MH patients. This led us to question ourselves on the reliability of the biomarkers found in case-control studies. Further analyses pinpointed four potential biomarkers for the development of MUO and CRC, although more investigation is necessary to confirm their role. This study can be added to the growing body of research aiming to include the GM in personalized medicine approaches.

## Supplementary Material

**Supplementary Figure 1.**
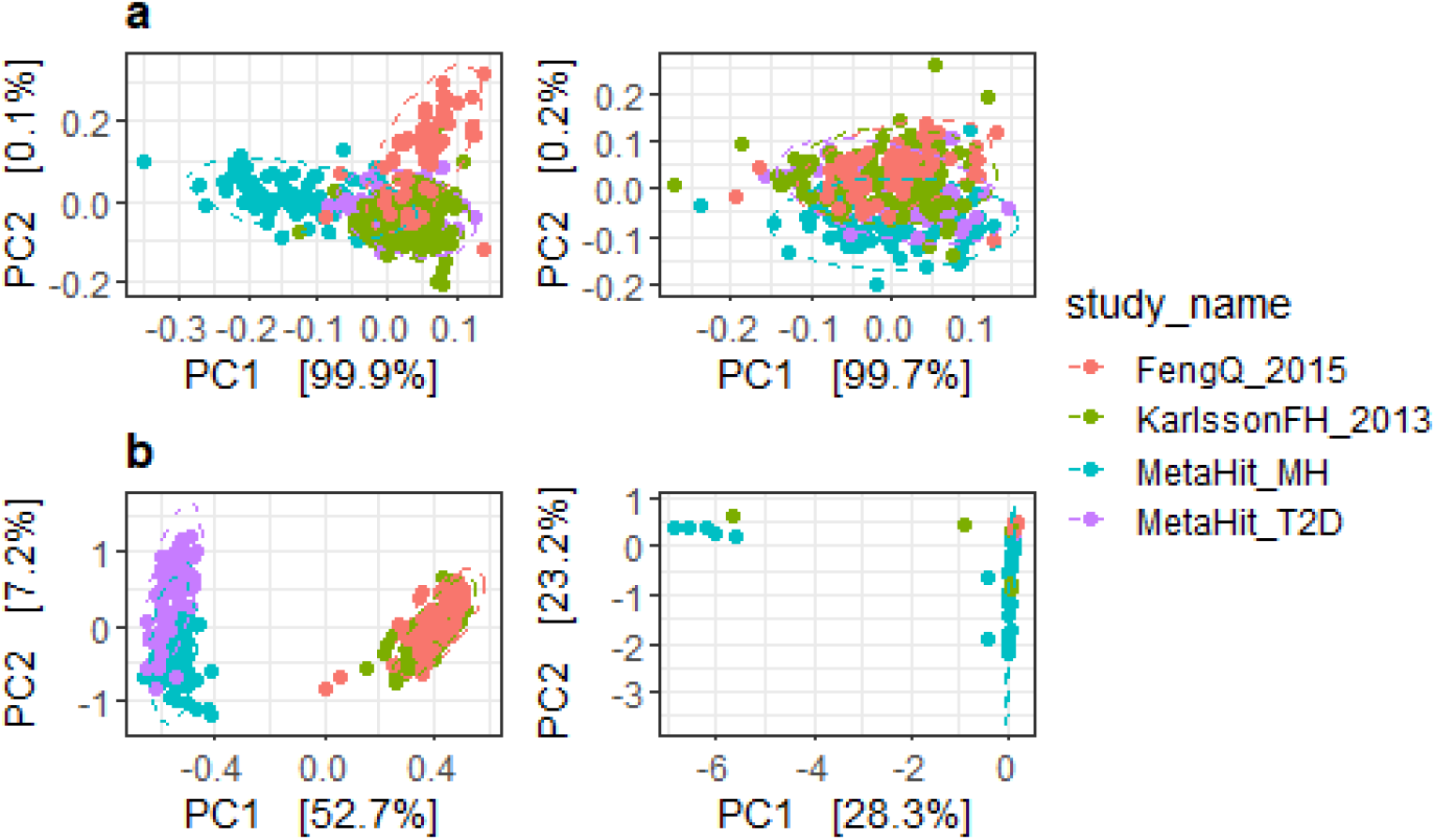
Batch effect correction with MMUPHin. Pathway abundance (**a**) and pathway coverage (**b**) files were imported to R and corrected using the *fit_adjust_function* from MMUPHin. This reduced the effect from the study variable from 65.18% to 9.16% (pathway abundances) and from 78.28% to 52.76% (pathway coverages), although it remained significant in both cases (*p* < 0.001, PERMANOVA with 999 permutations). Samples were ordinated via principal component analysis (PCA) and the two first components were plotted before (left) and after (right) correction. Ellipses represent 95% confidence intervals for each group.

**Supplementary Figure 2.**
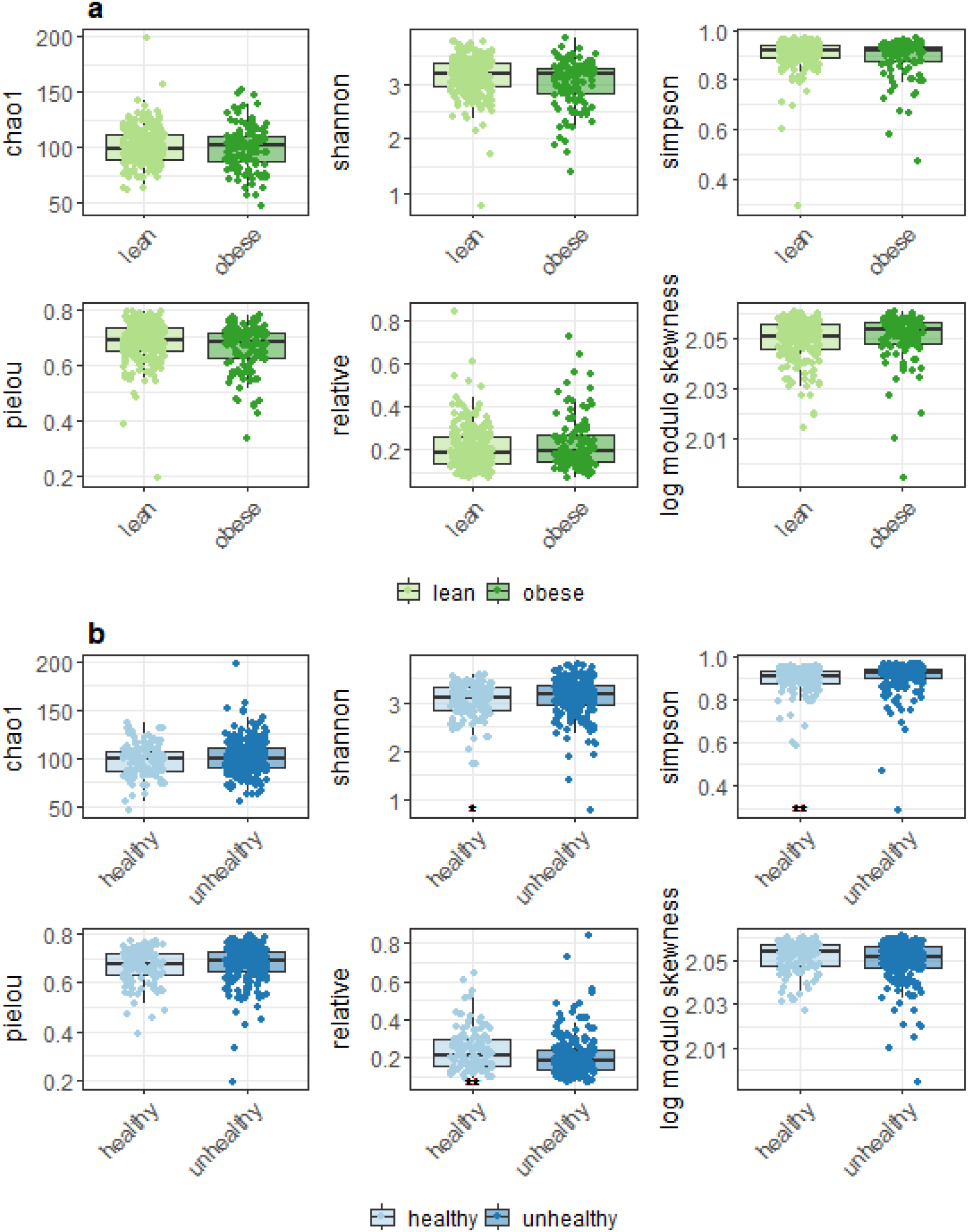
Alpha diversity analysis by obesity (**a**) or metabolic health status (**b**). Significances from the Wilcoxon test are indicated in the bottom left corner of the plot (healthy *vs*. unhealthy adjusted *p*-values are *p* = 0.0486, *p* = 0.0040 and *p* = 0.0024 for Shannon, Simpson and relative indices respectively).

**Supplementary Figure 3.**
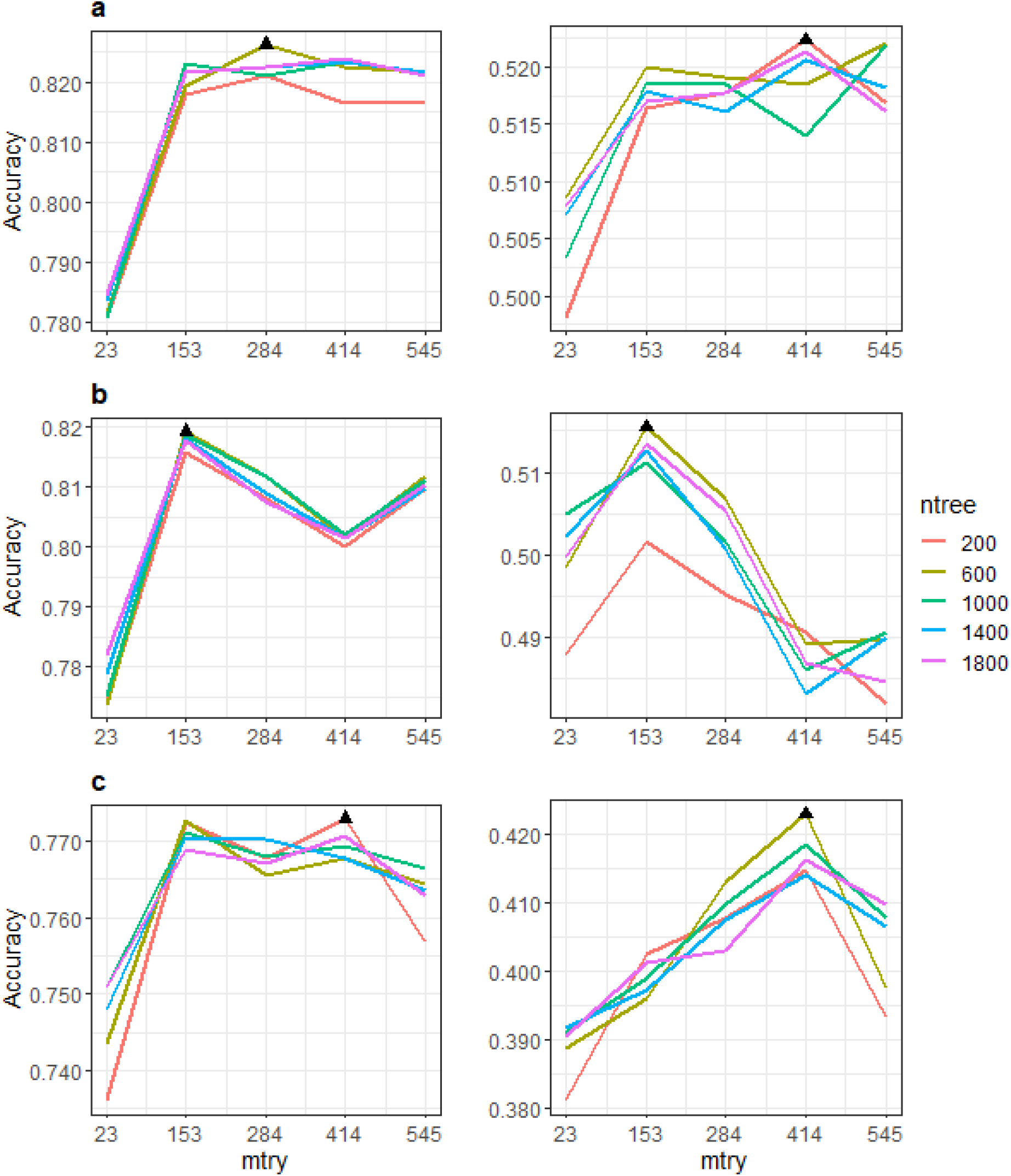
Hyperparameter tuning for models based on relative abundances together with age and sex, as well as body mass index in the binary models. Binary (left) and multiclass (right) models were trained without subsampling (**a**), upsampling (**b**) or downsampling (**c**) and with different *mtry* and *ntree* values. Black triangles show the hyperparameter combination reaching the highest accuracy for each model configuration.

**Supplementary Figure 4.**
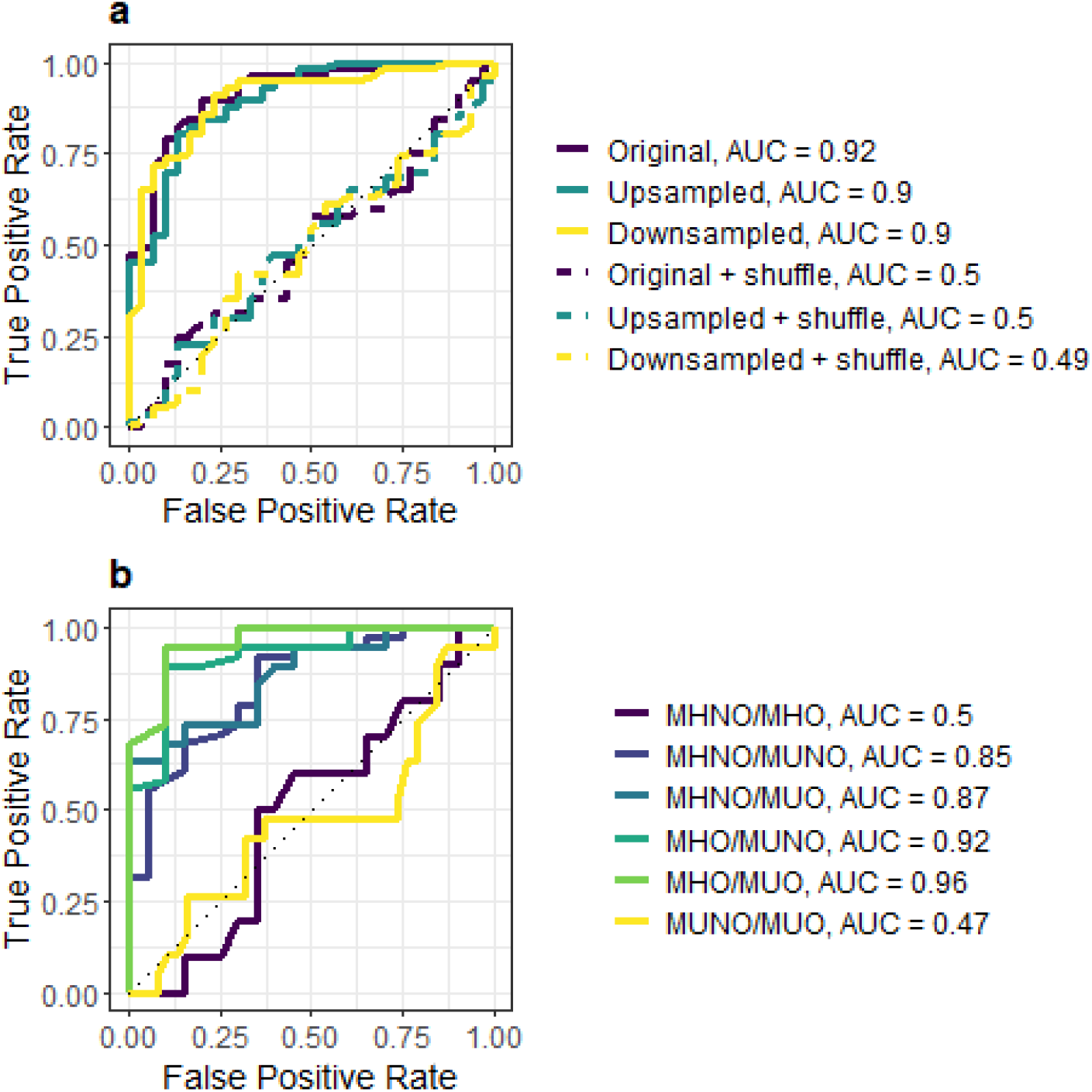
ROC curves for models based on relative abundances together with age and sex. **a)** The binary model with the best configuration for each subsampling technique as shown in Supp. Fig. 3 was used to predict class labels on the test set. Dashed lines show the ROC curves for a randomized version of the class labels. This model also used BMI information. **b)** ROC curves for every possible class combination are shown for the multiclass model with *ntree* = 200, *mtry* = 414 and without subsampling.

**Supplementary Figure 5.**
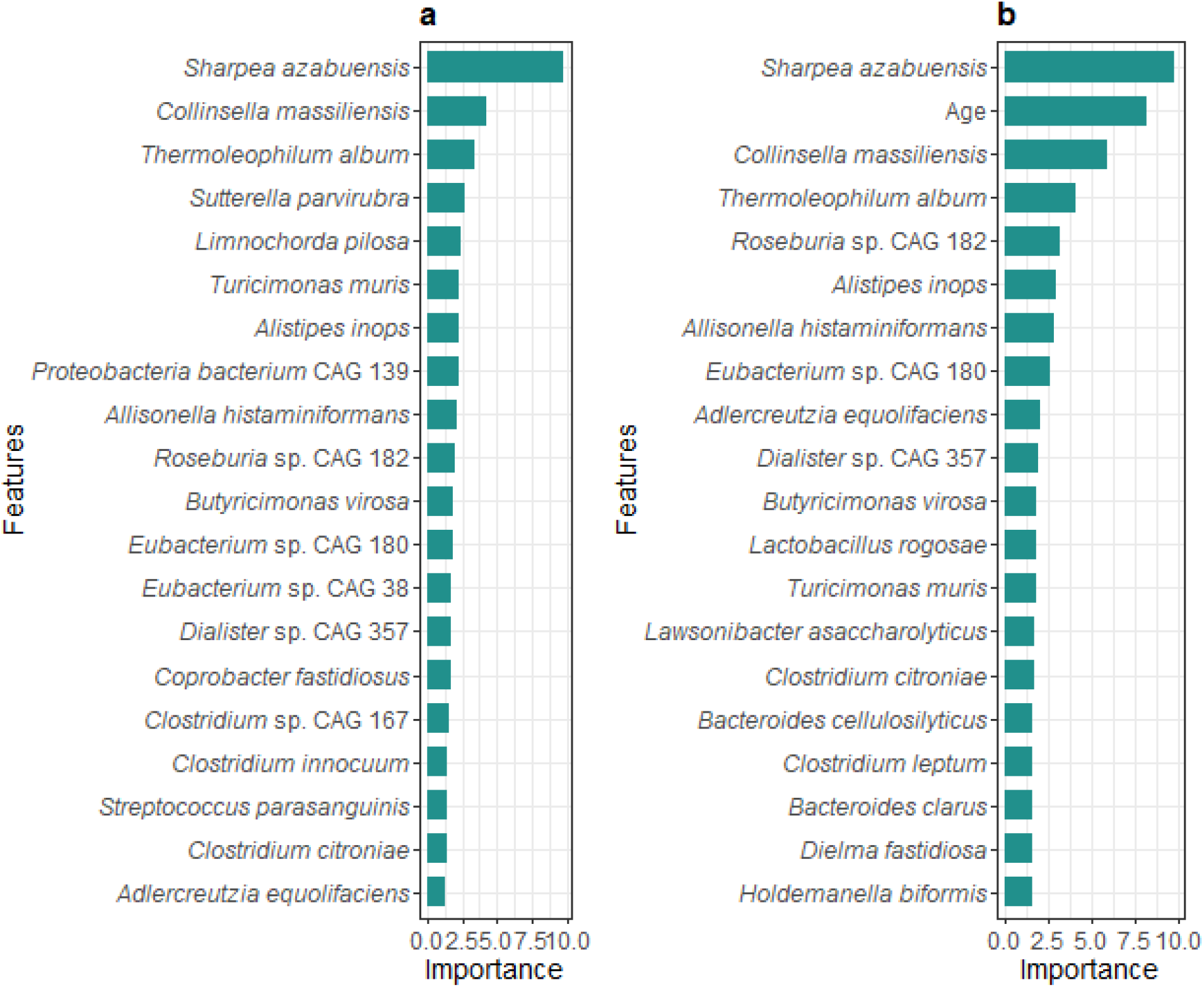
Top 20 features by importance in multiclass models. **a)** Top features in the best multiclass model based on metagenomic data alone (*ntree* = 600, *mtry* = 153). **b)** Top features in the best multiclass model including the subjects’ age and sex (*ntree* = 200, *mtry* = 414).

**Supplementary Figure 6.**
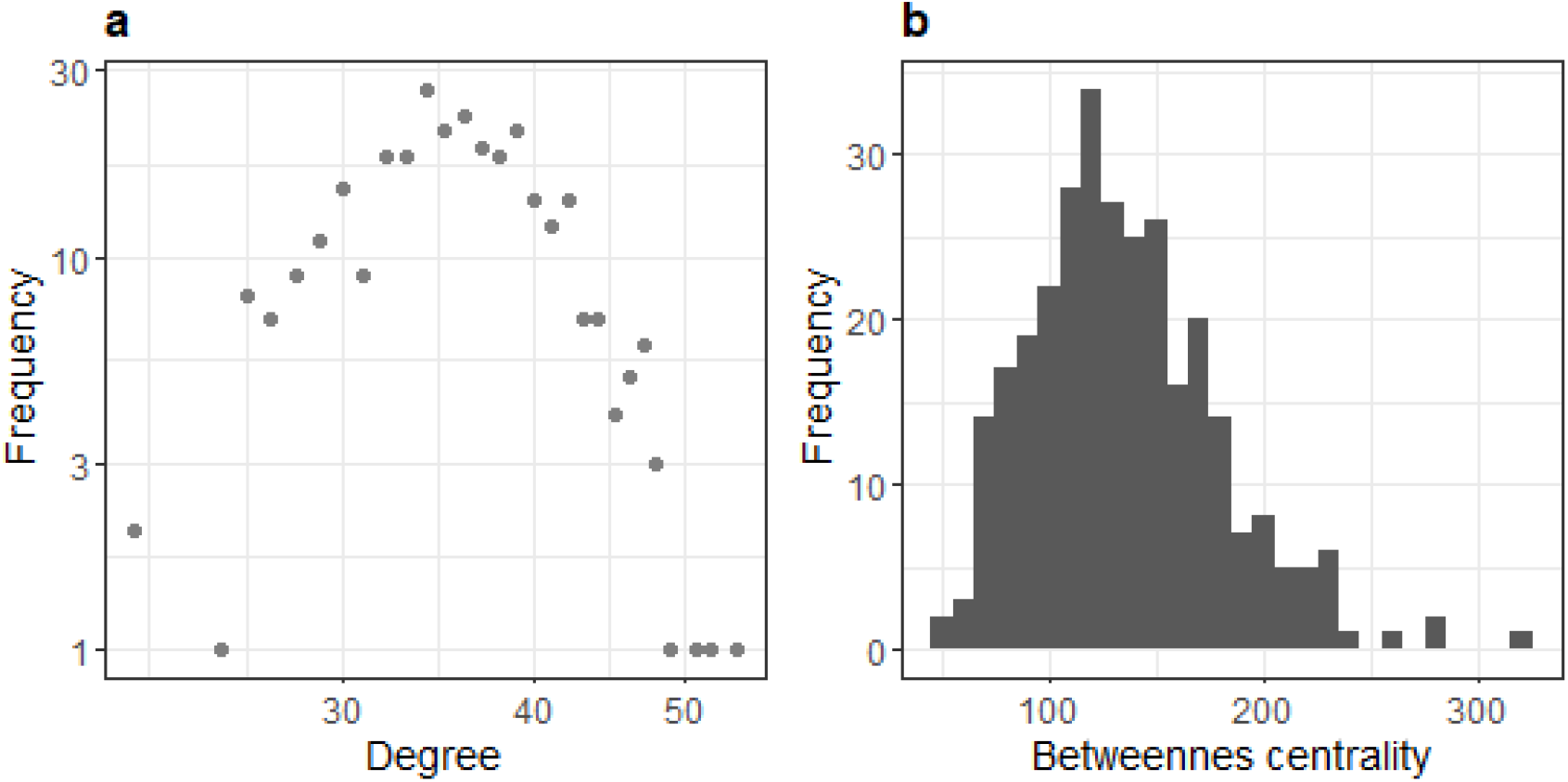
Analysis of a random graph equivalent to the patient similarity network. A random graph with 303 nodes and 63190 edges was created with the igraph function *erdos*.*renyi*.*game*. **a)** Node degree distribution (logarithmic scale). **b)** Node betweenness distribution.

**Supplementary Figure 7.**
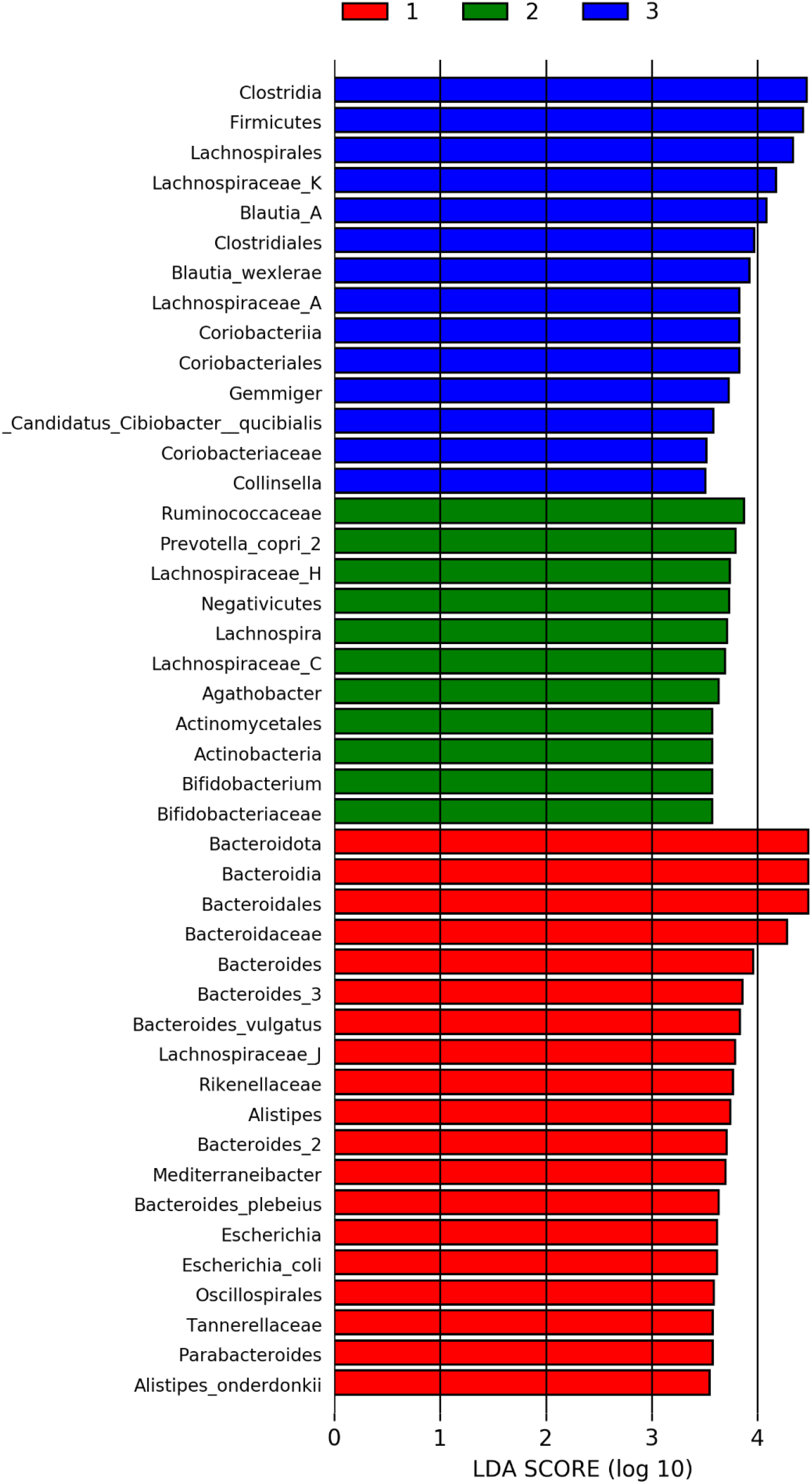
LEfSe of the ML4Microbiome COST Action patients. Barplot showing the LDA score of 43 features with *p* < 0.05 and LDA > 3.5. Red: CRC, green: control, blue: adenoma.

### List of abbreviations

16S rRNA: 16S ribosomal RNA gene
AUC: area under the curve
BMI: body mass index
CLR: centered log-ratio
CRC: colorectal cancer
CVD: cardiovascular disease
ENA: European Nucleotide Archive
FDR: false discovery rate
GM: gut microbiota
LDA: linear discriminant analysis
LEfSe: linear discriminant analysis of effect size
MetaHit: Metagenomics of the Human Intestinal Tract
MetS: metabolic syndrome
MH: metabolically healthy
MHNO: metabolically healthy non-obese
MHO: metabolically healthy obese
MSP: Metagenomic Species Pan-genome
MU: metabolically unhealthy
MUNO: metabolically unhealthy non-obese
MUO: metabolically unhealthy obese
PCA: principal component analysis
PCoA: principal coordinate analysis
PERMANOVA: permutational multivariate analysis of variance
RF: random forest
ROC: receiver operating characteristic
T2D: type 2 diabetes
UM: urolithin metabotype
WGS: whole-genome shotgun sequencing

## Declarations

### Ethics approval and consent to participate

Not applicable.

### Consent for publication

Not applicable.

### Author’s contributions

Conceptualisation, L.J.M.-Z., L.P.F., and E.C.d.S.P.; methodology, B.L.P., and L.J.M.-Z.; writing—original draft preparation, B.L.P. and L.J.M.-Z; writing—critical review and editing, E.C.d.S.P, L.P.F., A.R.d.M.; funding acquisition, A.R.d.M., L.J.M.-Z. and E.C.d.S.P. All authors have read and agreed to the published version of the manuscript.

### Corresponding authors

Correspondence to Enrique Carrillo de Santa Pau and Laura Judith Marcos-Zambrano.

